# Exploring thienothiadiazine dioxides as isosteric analogues of benzo- and pyridothiadiazine dioxides in the search of new AMPA and kainate receptor positive allosteric modulators

**DOI:** 10.1101/2023.11.02.565294

**Authors:** Pierre Francotte, Yasmin Bay, Eric Goffin, Thomas Colson, Cindy Lesenfants, Jerzy Dorosz, Saara Laulumaa, Pierre Fraikin, Pascal de Tullio, Caroline Beaufour, Iuliana Botez, Darryl S. Pickering, Karla Frydenvang, Laurence Danober, Anders Skov Kristensen, Jette Sandholm Kastrup, Bernard Pirotte

## Abstract

The synthesis and biological evaluation on AMPA and kainate receptors of new examples of 3,4-dihydro-2*H*-1,2,4-thieno[3,2-*e*]-1,2,4-thiadiazine 1,1-dioxides is described.

The introduction of a cyclopropyl chain instead of an ethyl chain at the 4-position of the thiadiazine ring was found to dramatically improve the potentiator activity on AMPA receptors, with compound **32** (BPAM395) expressing *in vitro* activity on AMPARs (EC2x = 0.24 µM) close to that of the reference 4-cyclopropyl-substituted benzothiadiazine dioxide **10** (BPAM344).

Interestingly, the 4-allyl-substituted thienothiadiazine dioxide **27** (BPAM307) emerged as the most promising compound on kainate receptors being a more effective potentiator than the 4-cyclopropyl-substituted thienothiadiazine dioxide **32** and supporting the view that the 4-allyl substitution of the thiadiazine ring could be more favorable than the 4-cyclopropyl substitution to induce marked activity on kainate receptors versus AMPA receptors.

The thieno-analogue **36** (BPAM279) of the clinically tested S18986 (**11**) was selected for *in vivo* evaluation in mice as a cognitive enhancer due to a safer profile than **32** after massive *per os* drug administration. Compound **36** was found to increase the cognition performance in mice at low doses (1 mg/kg) *per os* suggesting that the compound was well absorbed after oral administration and able to reach the central nervous system.

Finally, compound **32** was selected for co-crystallization with the GluA2-LBD (L504Y,N775S) and glutamate to examine the binding mode of thienothiadiazine dioxides within the allosteric binding site of the AMPA receptor. At the allosteric site, this compound established similar interactions as the previously reported BTD-type AMPA receptor modulators.

**Highlights:**

1. The study explored AMPA/kainate receptor PAMs belonging to thienothiadiazine dioxides
2. The 4-cyclopropyl-substituted compound **32** was the most potent AMPA receptor modulator
3. 4-Allyl substitution improved activity and selectivity for kainate receptors
4. The tricyclic compound **36** expressed cognitive improvement *in vivo* in mice
5. Compound **32** was co-crystallized with GluA2-LBD to examine receptor binding mode

**Graphical abstract:** 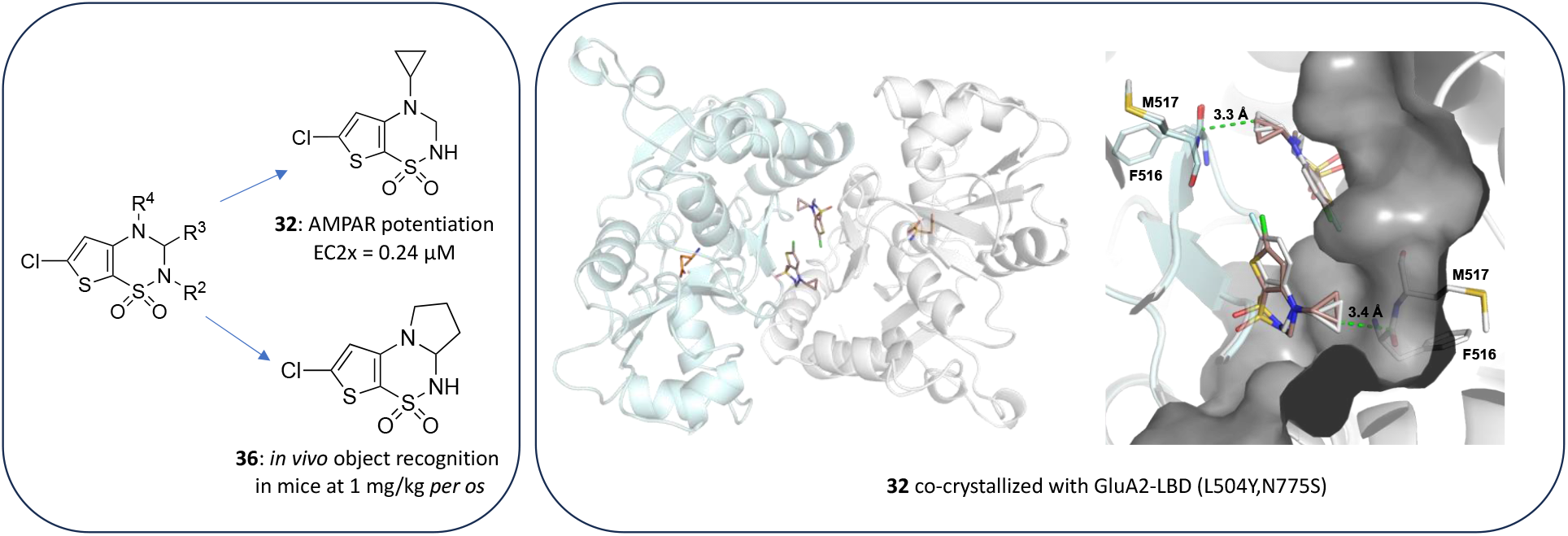

## 1. Introduction

L-Glutamate is the major excitatory neurotransmitter in the brain, interacting with ionotropic (ligand-gated ion channel, iGluRs) and metabotropic (G-protein coupled, mGluRs) receptors. The iGluR family is composed of four major subclasses of receptors: the *N*-methyl-*D*-aspartic acid (NMDA), the α-amino-3-hydroxy-5-methyl-4-isoxazolepropionic acid (AMPA), the kainic acid (KA) and the delta receptors [1, 2]. These receptors are essential for normal brain function but also play a role in many psychiatric and neurological disorders [3].

A decrease in AMPA receptor signaling is involved in Alzheimer’s disease, Parkinson’s disease, attention deficit hyperactivity disorder, depression, and schizophrenia [4]. Therefore, positive allosteric modulators (PAMs) of the AMPA receptors (AMPARs) are expected to become new drugs for the management of such neurodegenerative diseases [5–8]. Moreover, AMPAR PAMs stimulate the release of the brain-derived neurotrophic factor (BDNF), suggesting a therapeutic promise as neuroprotectants [9].

Kainate receptors (KARs) are also largely expressed in the brain and seem involved in several neurological and psychiatric disorders, such as epilepsy, schizophrenia, and depression [10–12]. However, little is known about the ability of small molecules to act as PAMs of the KARs (KAR PAMs).

AMPARs are tetrameric combinations of four subunits (GluA1-4) that arrange themselves in various stoichiometries, most of which being symmetric dimer-of-dimers of GluA2 and either GluA1, GluA3 or GluA4 [13,14]. KARs have very similar structural architecture but assemble from tetrameric combinations of five subunits (GluK1-5), forming homomeric or heteromeric receptors. Specifically, the GluK1-3 subunits can form functional homomeric and heteromeric receptors, whereas the two GluK4-5 subunits require co-assembly with GluK1-3 subunits to form functional heterotetrameric receptors [15].

In recent years, a great number of studies have focused on the identification and development of AMPAR PAMs belonging to distinct chemical classes, among which benzamides, 3,4-dihydro-2*H*-1,2,4-benzothiadiazine 1,1-dioxides (BTDs), and *N*-biaryl(cyclo)alkyl-2-propanesulfonamides [5,7,16]. Depending on their structure, these modulators were found to interact with distinct allosteric binding sites in the AMPAR [2,16–21].

Surprisingly, only a few examples of KAR PAMs have been described. To our knowledge, the BTD modulator BPAM344 (**10**, Figure 1) constitutes the first example of a small-molecule PAM of these receptors acting with different potency according to the receptor subtype [22]. Although BPAM344 is much more potent on AMPARs than on KARs, these results at least reveal that BTD-type compounds may serve as a starting point for discovering more potent KAR modulators after appropriate lead optimization.

**Figure 1.**
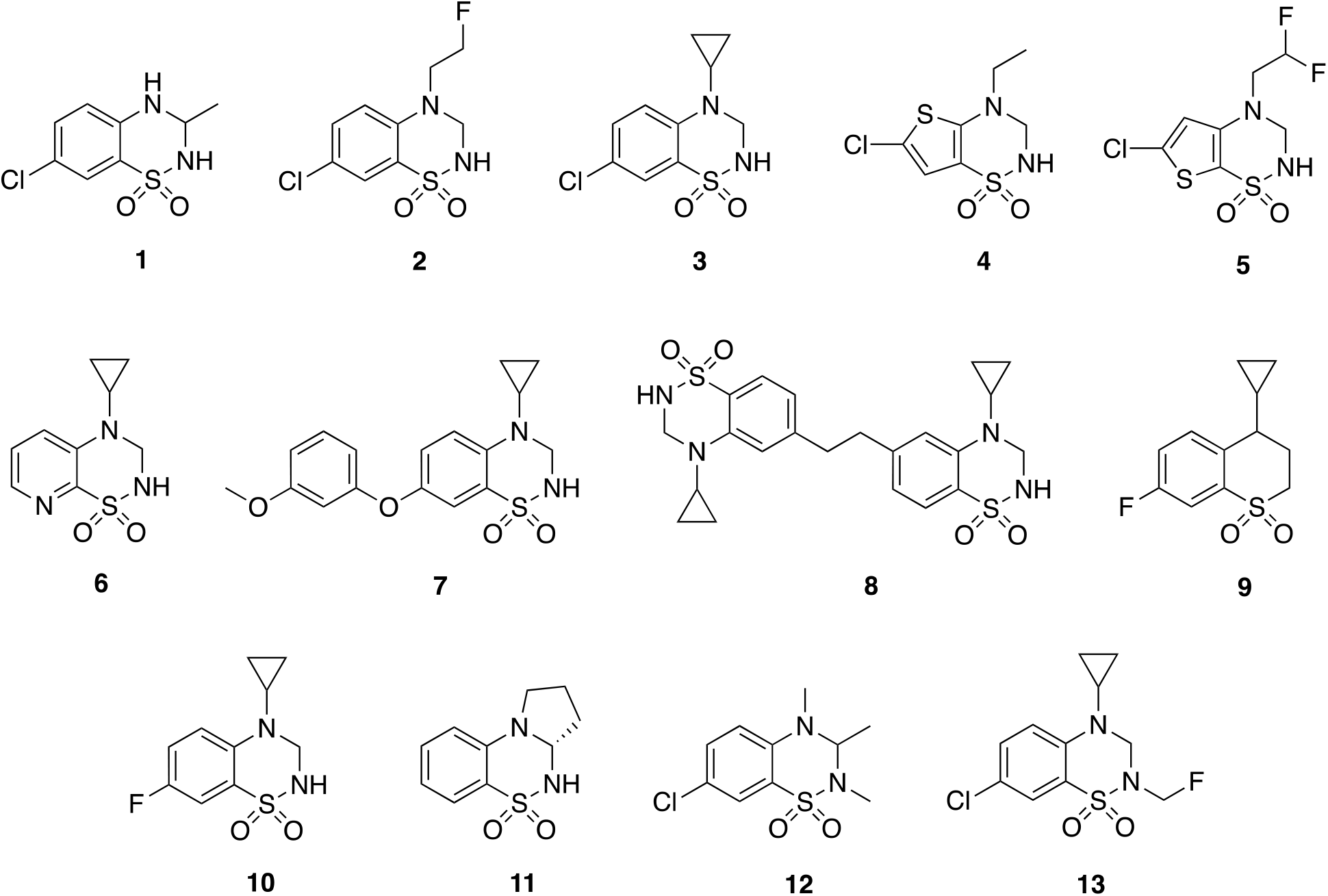
Examples of 3,4-dihydro-2*H*-1,2,4-benzothiadiazine 1,1-dioxides (**1**-**3**, **7**, **8**, **10-13**), one example of 3,4-dihydro-2*H*-thieno[2,3-*e*]-1,2,4-thiadiazine 1,1-dioxide (**4**), one example of 3,4-dihydro-2*H*-thieno[3,2-*e*]-1,2,4-thiadiazine 1,1-dioxide (**5**), one example of 3,4-dihydro-2*H*-pyrido[3,2-*e*]-1,2,4-thiadiazine 1,1-dioxide (**6**), and one example of thiochroman 1,1-dioxide (**9**) reported as AMPAR PAMs.

We have recently focused on developing new AMPAR PAMs belonging to the BTD-type compounds starting from the structure pattern of the orally active AMPAR potentiator IDRA 21 (**1**, Figure 1) [23–35]. New orally active BTD compounds such as **2** and **3** (Figure 1) were reported to express improved *in vitro* properties and *in vivo* efficacy [25, 29]. The isosteric replacement of the benzene ring of BTDs by a thiophene (i.e., **4** and **5**, Figure 1) or a pyridine ring (i.e., **6**, Figure 1) was also explored with success [27,29]. More recently, 7-phenoxy-substituted BTDs such as **7** (Figure 1) [33] and dimers such as **8** (Figure 1) [34] have been shown to be particularly potent as AMPAR modulators, potentiating the effect of glutamate on these receptors in the nanomolar range [33,34]. Based on the isosteric replacement concept, thiochroman 1,1-dioxides like **9**, which is the structural analogue of the potent compound **10**, were also found to express a noticeable potentiator activity on the AMPARs [35]. Finally, examples of BTDs bearing an alkyl substituent at the 2- and/or the 3-position(s) were also explored, providing interesting potentiators of the AMPARs [31]. The BTD compound S18986 (**11**) (Figure 1) from Servier is an example of a ring-closed 3,4-dialkyl-disubstituted BTD that reached clinical trials [36,37]. The 2,3,4-trimethyl-substituted BTD **12** (Figure 1) was surprisingly found to be a potent AMPAR PAM *in vitro,* while the corresponding 2,3- and 3,4-dimethyl-substituted analogues were substantially less active [31]. Recently, compound **13** bearing a fluoromethyl group at the 2-position was found to exert better *in vivo* activity (improvement of cognitive functions) after oral administration in mice compared to its 2-unsubstituted analogue **3** [38].

In a continuous effort to discover more promising AMPAR and KAR modulators, the present study explored new examples of variously substituted 6-chloro-3,4-dihydro-2*H*-thieno[3,2-*e*]-1,2,4-thiadiazine 1,1-dioxides. More specifically, this study considered compounds bearing new kinds of alkyl groups at the 4-position (cyclopropyl, allyl) and examples of 2,4-dialkyl-/2,3,4-trialkyl-substituted compounds, including ring-closed analogues of **11** as well as the 2-methyl- and 2,3-dimethyl-substituted analogues of compound **5** previously reported as the most potent thieno[3,2-*e*]-1,2,4-thiadiazine 1,1-dioxide [27]. Particular attention was paid to the introduction of a cyclopropyl chain instead of an ethyl chain at the 4-position of the thiadiazine ring since this structural modification has been found to dramatically improve the potentiator activity at AMPARs for previously reported benzo- and pyridothiadiazine dioxides (Figure 2) [28,29]. Moreover, the synthesis and biological evaluation of tricyclic compounds, corresponding to thieno-analogues of the clinically tested S18986 (**11**), were also examined (Figure 2).

**Figure 2.**
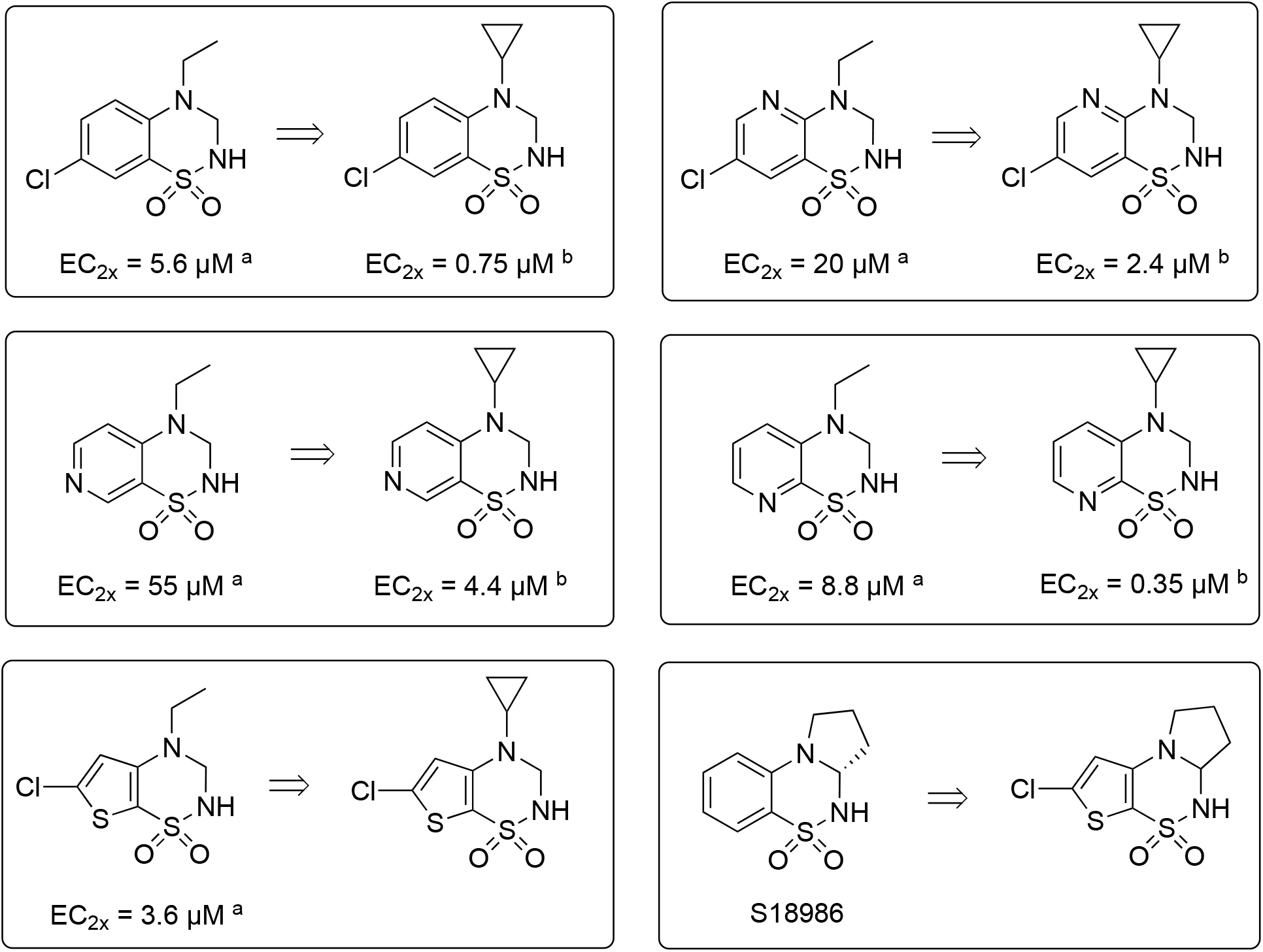
At the top (the first four boxes): replacement of the ethyl chain at the 4-position of the heterocycle with a cyclopropyl chain in different series of ring-fused thiadiazine dioxides leading to a strong improvement of the *in vitro* potentiator activity on AMPARs (^a^ EC2x values established by voltage clamp on *Xenopus* oocytes expressing the AMPARs; ^b^ EC2x values established by fluorescence on primary cultures of neurons from rat embryonic cortex. At the bottom (the last two boxes): new developments projected in this work.

## 2. Results and discussion

### 2.1. Chemistry

The synthetic pathways used to prepare the 6-chloro-3,4-dihydro-2*H*-thieno[3,2-*e*]-1,2,4-thiadiazine 1,1-dioxides reported here are illustrated in Schemes 1-5.

The ring closure reaction of **14** (obtained according to [27]) with acetaldehyde in the presence of a catalytic amount of camphorsulfonic acid provided the unstable 3,4-dialkyl-substituted intermediate **15**. The latter was immediately converted into the stable 2-methyl-substituted analogue **16** after alkylation with methyl iodide in acetonitrile in the presence of potassium carbonate (Scheme 1).

Starting from the *o*-aminothiophenesulfonamide **17** hydrochloride synthesized as previously described [27], acylation of the primary amine function using difluoroacetyl chloride provided intermediate **18** (Scheme 2). The latter reacted with lithium aluminum hydride in diethyl ether to convert the secondary amide **18** into the corresponding secondary amine **19**. Ring closure reaction by heating compound **19** in formic acid led to the expected 4*H*-thieno-1,2,4-thiadiazine 1,1-dioxide **20**. Saturation of the N=C double bond of **20** after reaction with sodium borohydride in isopropanol provided intermediate **21**, which was alkylated by methyl iodide in acetonitrile in the presence of potassium carbonate to give the target compound **22** (Scheme 2). The corresponding compound bearing a methyl group at the 3-position was obtained after ring closure reaction of **19** by means of acetaldehyde under acidic catalysis, leading to the unstable intermediate **23**, which was immediately alkylated at the 2-position after reaction with methyl iodide in acetonitrile in the presence of potassium carbonate to provide the target compound **24** (Scheme 2).

The 4-allyl-substituted compounds **27** and **28** were obtained as described in Scheme 3. Starting from intermediate **17**, ring closure reaction with triethyl orthoformate led to 6-chloro-4*H*-thieno[3,2-*e*]-1,2,4-thiadiazine 1,1-dioxide **25**. Alkylation in the 4-position of this heterocycle by means of allyl bromide led to intermediate **26**, which was reduced with sodium borohydride to provide the target compound **27**. The latter was alkylated by methyl iodide in acetonitrile in the presence of potassium carbonate to give the target compound **28** (Scheme 3).

The *o*-aminothiophenesulfonamide **17** hydrochloride was also used to synthesize the target compound **32** (Scheme 4). Although introduction of the cyclopropyl moiety at the 4-position of intermediate **25** by direct alkylation in the presence of cyclopropyl bromide was not successful, we used an alternative route using (1-ethoxycyclopropoxy)trimethylsilane in methanol and acetic acid. This reaction led to the 1-methoxycyclopropylamino-substituted intermediate **29** instead of the expected 1-ethoxycyclopropylamino-substituted compound due to a probable exchange of the ethoxy group with the methoxy group from the reaction solvent methanol. Intermediate **30** was obtained after reaction of **29** with sodium borohydride and boron trifluoride diethyl etherate in THF. Ring closure reaction on **30** in the presence of triethyl orthoformate led to intermediate **31**, which reacted with sodium borohydride in isopropanol to provide the target compound **32** (Scheme 4).

Lastly, the tricyclic compounds **36** and **37** were prepared as described in Scheme 5. The starting compound **17** hydrochloride reacted with 4-chlorobutyryl chloride in dioxane to form the corresponding amide **33**. By heating the latter in 1N NaOH for 12 h, a ring closure reaction formed the corresponding pyrrolidinone intermediate **34**. The fusion of this intermediate at 200-210 °C produced a second ring closure to give access to the tricyclic compound **35**. Saturation of the N=C double bond of **35** by using sodium borohydride in isopropanol led to the target compound **36**. Methylation of the nitrogen atom of the sulfonamide function of **36** by means of methyl iodide in the presence of potassium carbonate provided the second target compound **37** (Scheme 5).

### 2.2. In vitro evaluation

The new thieno[3,2-*e*]-1,2,4-thiadiazine 1,1-dioxides were first evaluated as AMPAR potentiators using a previously described *in vitro* membrane potential fluorescence-based assay (FlipR or FDSS) for activity at native AMPARs in rat cortical primary cell cultures [25]. For each compound, the EC2x value was determined, corresponding to the concentration of PAM responsible for a 2-fold increase of the amplitude of the current induced by AMPA at 300 μM (Table 1). For evaluation at recombinant AMPAR and KAR subtypes fluorescence-based Ca^2+^-influx assays were used (Figure 3A-C and Table 1). Compound ability to potentiate glutamate-evoked AMPAR activity was tested at homomeric GluA2 receptors stably expressed in HEK293-GT cells. Notably, GluA2 receptors contain a glutamine residue at a site in the ion channel (denoted the Q/R site) that in native receptors is changed to an arginine (R) as a result of RNA editing, which renders the channel Ca^2+^-impermeable [39]. Also, GluA2 is exposed to alternative mRNA splicing, resulting in two isoforms known as flip (i) or flop (o) [40] that have different effects on PAM potency and effect. For the present assay, the non-edited (Q) version of the flip isoform of GluA2 (GluA2(*Q*)_i_) was used. The compounds were first tested for the ability to potentiate AMPAR activity at a concentration of 100 µM. Specifically, GluA2(*Q*)_i_ mediated Ca^2+^ influx was evoked by a saturating concentration of L-glutamate (100 µM) in the absence and presence of the test compounds and compared to the effect of the reference compound **10** (BPAM344) at 100 µM (Figure 3A). Dose-response curves were established for compounds displaying potentiation to determine the EC_50_ value for the potentiating effect at GluA2(*Q*)_i_ (Figure 3B). Compound **37** could not reach concentrations high enough to construct a dose-response curve due to a solubility issue in the assay buffer (maximum tested concentration 1000 µM). Moreover, the four most potent compounds on AMPARs (**16**, **24**, **27**, **32**) and the reference compound **10** were tested at homomeric GluK1(*Q*)_1b_ and GluK2(*Q*)_2a_ receptors using 100 µM KA as agonist (Figure 3C).

**Table 1.** Effects of 4-alkyl-, 3,4-dialkyl- and 2,3,4-trialkyl-substituted 6-chloro-3,4-dihydro-2*H*-thieno[3,2-*e*]-1,2,4-thiadiazine 1,1-dioxides on the fluorescence induced by 300 μM AMPA on primary cultures of neurons from rat embryonic cortex (FlipR/FDSS) and on the calcium influx induced by 100 µM glutamate on HEK293 dells stably expressing the GluA2(*Q*)_i_ subunit.

| Compd. | R<sup>2</sup> | R<sup>3</sup> | R<sup>4</sup> | EC2x<br>(FlipR<sup>a</sup>/FDSS<sup>b</sup>) (μM) | Ca<sup>2+</sup> influx<br>EC<sub>50</sub> (μM)<sup>c</sup> |
| --- | --- | --- | --- | --- | --- |
| 16 | -CH<sub>3</sub> | -CH<sub>3</sub> | -CH<sub>2</sub>CH<sub>3</sub> | 6.4 [3.2; 12.9] (2)<sup>a</sup> | 24.6 ± 6.6 (3) |
| 5<sup>d</sup> | -H | -H | -CH<sub>2</sub>CHF<sub>2</sub> | 2.9 ± 0.1 (3)<sup>a,d</sup> | n.d. |
| 22 | -CH<sub>3</sub> | -H | -CH<sub>2</sub>CHF<sub>2</sub> | 24.5 [14.7; 34.3] (2)<sup>b</sup> | 39.9 ± 8.6 (3) |
| 24 | -CH<sub>3</sub> | -CH<sub>3</sub> | -CH<sub>2</sub>CHF<sub>2</sub> | 1.2 [0.7; 1.7] (2)<sup>b</sup> | 31.5 ± 11.3 (3) |
| 27 | -H | -H | -CH<sub>2</sub>CH=CH<sub>2</sub> | 18.5 [12.1; 29.4] (3)<sup>a</sup> | 26.9 ± 2.4 (3) |
| 28 | -CH<sub>3</sub> | -H | -CH<sub>2</sub>CH=CH<sub>2</sub> | 80.4 [47.9; 135.0] (3)<sup>a</sup> | 76.5 ± 7.7 (3) |
| 32 | -H | -H | -CH(CH<sub>2</sub>)<sub>2</sub> | 0.24 [0.15; 0.34] (2)<sup>b</sup> | 4.7 ± 0.9 (3) |
| 36<sup>e</sup> | -H | -CH<sub>2</sub>CH<sub>2</sub>CH<sub>2</sub>- | | 33.5 [29.3; 38.2] (3)<sup>a</sup> | 40.2 ± 8.0 (3) |
| 37 | -CH<sub>3</sub> | -CH<sub>2</sub>CH<sub>2</sub>CH<sub>2</sub>- |  | 40.1 [37.6; 42.7] (2)<sup>a</sup> | > 100 (3) |
| 10<sup>d</sup> |  |  |  | 0.26 [0.15; 0.47] (3)<sup>b,d</sup> | n.d. |

<sup>a</sup> EC2x: concentration of modulator giving a 2-fold increase of the fluorescence induced by AMPA (300 μM) with the FlipR method on rat primary brain cultures (n = 2–3); results are expressed as the geometric mean with its 95% confidence intervals in brackets. <sup>b</sup> EC2x values obtained by the FDSS fluorescence assay, a variant of the FlipR fluorescence assay using another plate reader. <sup>c</sup> EC<sub>50</sub>: AMPAR potentiator concentration responsible for 50% of the maximal effect (mean ± SEM (n = 3)). <sup>d</sup> Published compounds and results [27, 38]; for **5** the result is given as the mean ± SEM (n) [27]. <sup>e</sup> Compound **36** tested on *Xenopus* oocytes (magnitude of the current induced by AMPA (10 μM) in *Xenopus* oocytes injected with rat cortex mRNA according to [24] expressed an EC2x value of 15.4 ± 2.6 μM and an EC<sub>50</sub> value of 48.6 ± 1.5 μM.

**Figure 3.**
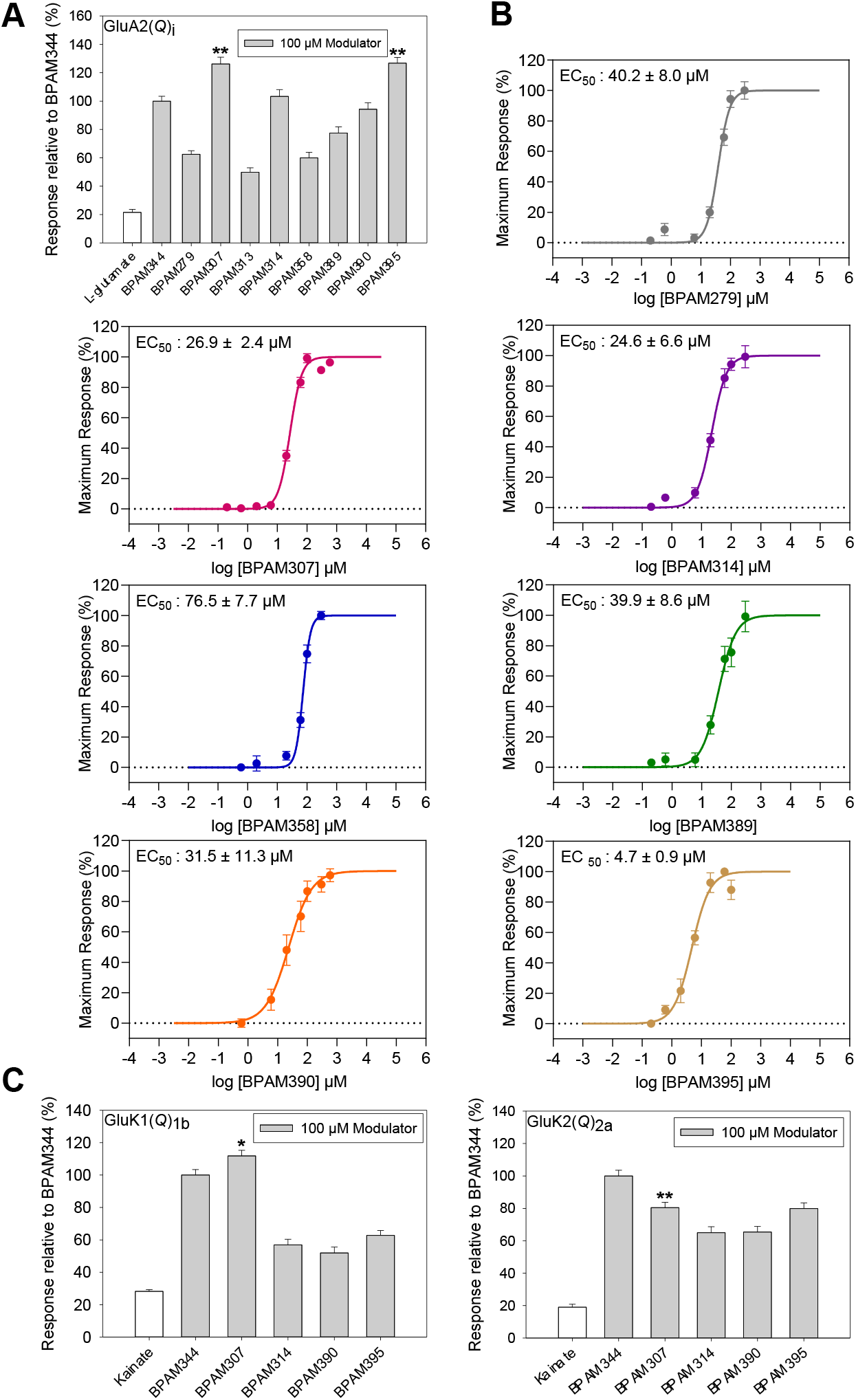
Pharmacological characterization of the new compounds using calcium-sensitive fluorescence-based assays (BPAM344 = **10**; BPAM279 = **36**; BPAM307 = **27**; BPAM313 = **37**; BPAM314 = **16**; BPAM358 = **28**; BPAM389 = **22**; BPAM390 = **24**; BPAM395 = **32**). (**A**) Evaluation of modulators at GluA2(*Q*)_i_ in the presence of 100 µM glutamate. Recorded RFU was normalized to the response of BPAM344 (**10**) and shown as mean ± SEM from at least three independent experiments. (**B**) Dose-response curves of the modulators at GluA2(*Q*)_i_ in the presence of 100 µM glutamate. Each data point represents the mean ± SEM from triplicate wells (n = 9-12). The dose-response curves are generated from pooled data (at least three experiments). EC_50_ values are shown as mean ± SEM. (**C**) Evaluation of modulators at GluK1(*Q*)_1b_ and GluK2(*Q*)_2a_ in the presence of 100 µM KA.

Table 1 indicates that, regarding the EC2x values, the introduction of a methyl group at the 2,3-positions was responsible for the preservation of a marked potentiator activity on AMPARs (compare **24** versus **5**), while the 2-monomethylation appeared to be less favorable (compare **24** versus **22**; **28** versus **27**; **37** versus **36**). The EC_50_ values obtained in the second biological assay evolved in the same way, although the EC_50_ values were generally found to be higher than the EC2x values.

Interestingly, the introduction of a cyclopropyl group at the 4-position of the heterocycle markedly improved the potentiator activity on the AMPARs (see compound **32**: EC2x = 0.24 µM; EC_50_ = 4.7 µM). This observation agrees with previous results obtained in the benzo- and pyridothiadiazine dioxide series (Figure 2).

Compound **36**, the 6-chlorothieno-analogue of the tricyclic BTD compound **11** (S18986), expressed a moderate *in vitro* activity on AMPARs although comparable to the latter (**11**: *Xenopus* oocytes: EC2x = 25 µM [36]; FDSS: 40 µM, unpublished result).

Tested at 100 µM on GluA2, compounds **27** and **32** were found to have higher efficacy (∼126%) than the reference compound **10** normalized to 100% (P-value <0.001, Student’s t-test, SigmaPlot, ** in histogram, Figure 3A), whereas the remaining compounds displayed lower efficacy except compound **16** with same efficacy.

Compound **27** tested at 100 µM expressed higher efficacy (112%) than compound **10** normalized at 100% at GluK1(*Q*)_1b_ (P-value = 0.019, Student’s t-test, SigmaPlot, * in histogram, Figure 3C), whereas **16**, **24**, and **32** were found to have lower efficacy (50-60% relative to **10**). At GluK2(*Q*)_2a_, all four modulators expressed lower efficacy (**16** and **24** ∼60% and **27** and **32** ∼80%, P-value <0.001, Student’s t-test, SigmaPlot, ** in histogram, Figure 3C) relative to **10**. From these results, compound **27** (BPAM307) emerged as the best thienothiadiazine dioxide on KARs, supporting the view that the allyl side chain at the 4-position of the thiadiazine ring could be more favorable than the cyclopropyl chain to induce marked activity and selectivity for the KARs versus AMPARs. In fact, compound **32,** bearing a cyclopropyl chain at the 4-position of the heterocycle, although very potent on AMPARs, was found to express a lower potentiator activity on KARs.

### 2.3. In vivo evaluation

The *in vitro* biological data identified compound **32** as a possible candidate for further investigations and development as a new drug acting on AMPARs. However, to select the best candidate, we examined the safety profile of the new compounds *in vivo* by determining their possible pro-convulsant effect or induction of hypothermia resulting from a massive *per os* drug administration in NMRI mice (30 mg/kg and 100 mg/kg). Indeed, a major concern with AMPAR PAMs is seizure induction at high doses resulting from an excessive stimulation of the glutamatergic neurotransmission in the CNS [41,42]. Unfortunately, compound **32** induced hypothermia at 30 mg/kg *per os* in NMRI mice, while compound **36** was remarkably safe up to 100 mg/kg (no hypothermia and convulsions; see Figure 4).

**Figure 4.**
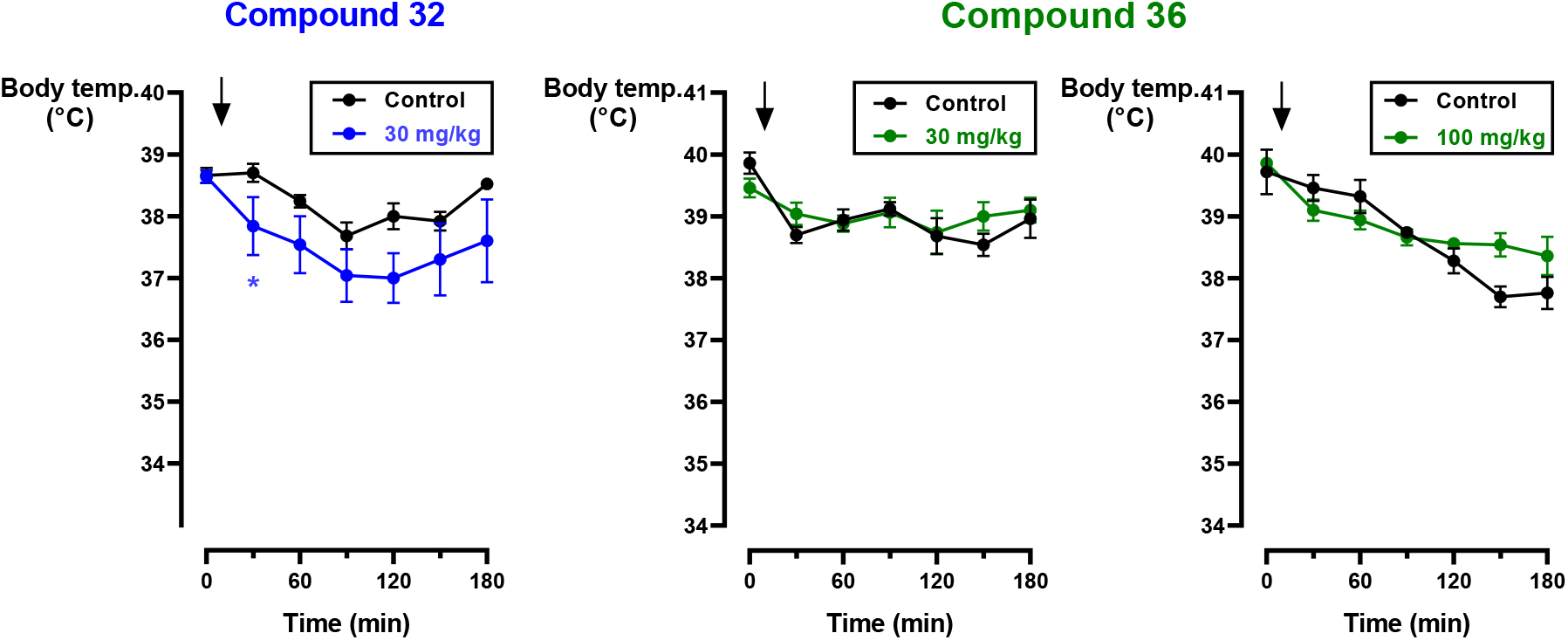
Effect of compound **32** and compound **36** after *per os* administration (30 mg/kg and 100 mg/kg) on the body temperature In NMRI mice. Data is expressed as means ± SEM (n = 5/group) (*: p<0.05 vs. vehicle control group).

Considering the potential benefit of **36** over the other thienothiadiazine dioxides, we decided to investigate the effects of **36** in two *in vivo* experiments.

We first examined the effects of compound **36** on long-term potentiation (LTP) of the post-synaptic responses induced *in vivo* by tetanic stimulation (Figure 5). LTP is thought to represent a synaptic plasticity mechanism that underlies learning and memory processes [43,44]. Specifically, one hour after the intraperitoneal administration of **36** or vehicle, LTP in the dentate gyrus of the hippocampus in anesthetized rats was induced by a brief high-frequency stimulation of the perforant path, and the subsequent excitatory postsynaptic field potentials (EPSfPs) were then recorded during 3 h. Compared to vehicle, **36** was found to increase the amplitude of the initial post-synaptic response prior to the tetanic stimulation but also the duration of the LTP at a dose of 30 mg/kg (Figure 5). The efficacy observed with **36** was comparable to that obtained with previously described BTDs, albeit at higher doses [25]. However, this effect demonstrated that the compound was able to cross the blood-brain barrier and to reach the CNS, supporting the view that it may exert a cognition-enhancing effect.

**Figure 5.**
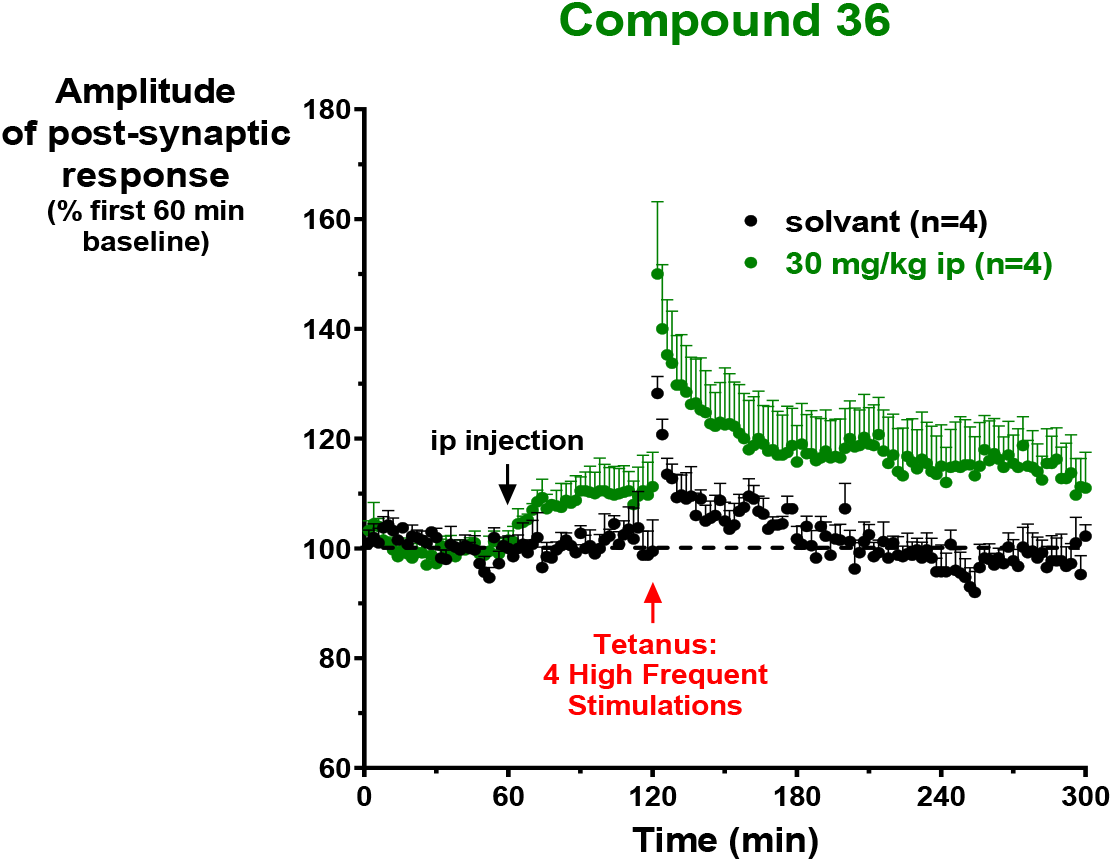
Post-synaptic response induced *in vivo* in the anesthetized rat. Effect of compound **36** (30 mg/kg i.p.) on the potentiation of the synaptic response evoked in the dentate gyrus of the hippocampus by a high-frequency stimulation (HFS) delivered in the perforant path. In each rat, synaptic responses were averaged over four successive stimulations (every 2 min) and normalized to the average amplitude of synaptic responses obtained during the first 60 min baseline period before the injection of the compounds (control value taken as 100%). Results are expressed as mean ± SEM.

To investigate potential improvement of cognitive performance *in vivo*, the effect of **36** was evaluated in an object recognition test in CD1 mice (Figure 6). The three-session test performed, considered as a paradigm for episodic memory in rodents, was based on that animals remembering a familiar object seen in the previous session spend less time exploring it compared to exploring a new object [45]. According to Figure 6, oral administration of **36** one hour before the three sessions increased the cognition performance of mice at doses as low as 1 mg/kg. This effect on the object recognition test in mice confirmed the interest of **36** as a cognitive-enhancing drug and suggests that the compound is absorbed in mice after oral administration and reach the CNS. The percentage of increase of the exploration of the new object versus the familiar one was +35 % for vehicle and +82 % and +69 % for compound 36 at 1 and 3 mg/kg, respectively (Figure 6). When considering the difference of exploration between New and Familiar objects (New - Fam), a trend of increase was thus noticed. However, the difference did not reach statistical significance due to high variability in the vehicle control group (ANOVA p>0.05).

**Figure 6:**
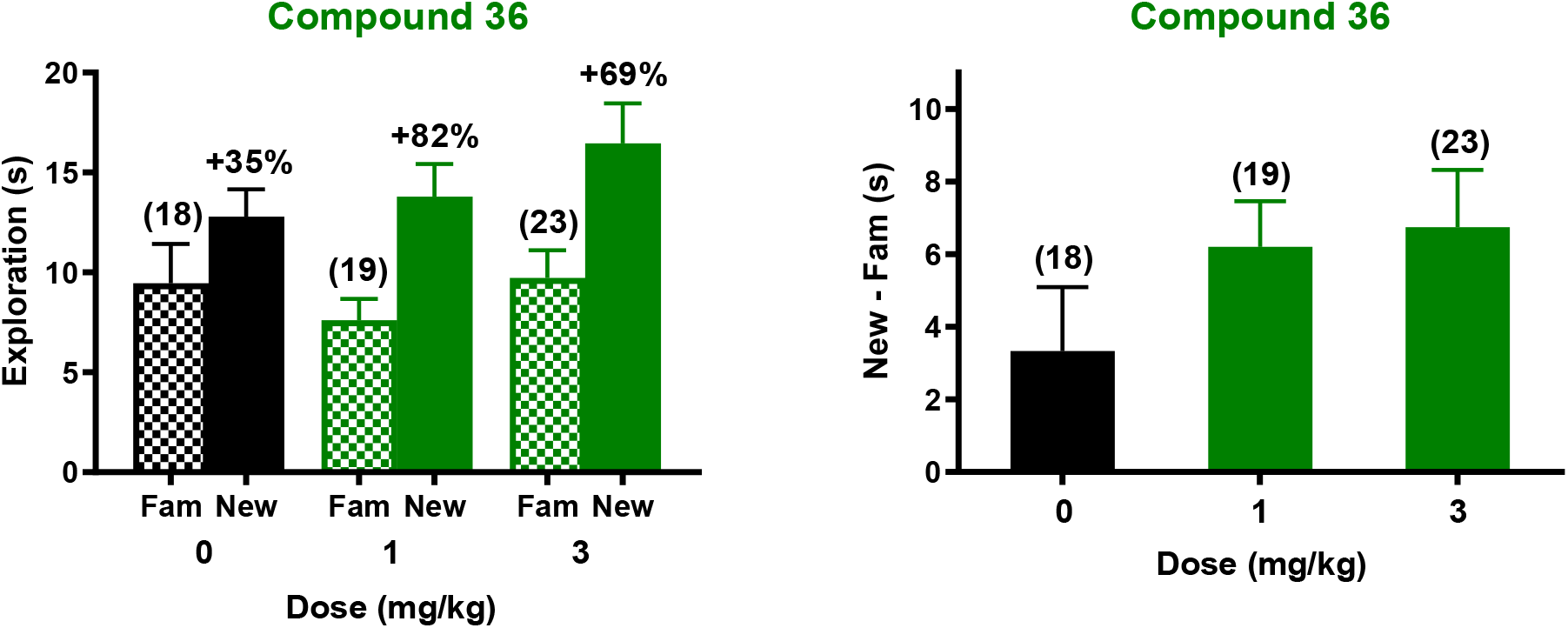
Effect of Compound **36** (1 and 3 mg/kg, *per os*) on the exploration time spent on either the Familiar or the New object during the recognition phase in the object recognition task in CD1 mice. During the recognition phase, compound **36** at either 1 or 3 mg/kg PO increased the duration of exploration of the new object *versus* the familiar one, whereas there is no difference in the vehicle-treated group (on the left). The number in brackets refers to the number of animals. The % of effect was +35 %, +82 % and +69 % for vehicle, 1 and 3 mg/kg, respectively.

### 2.4. Structure of the ligand-binding domain of GluA2 with glutamate and compound **32**

The ligand-binding domain of rat GluA2, containing the mutations L504Y and N775S (GluA2-LBD) was crystallized with glutamate and the most potent AMPAR modulator within the series, compound **32**. The L504Y mutation creates a dimeric GluA2-LBD in solution [46], whereas the N775S mutation makes it a flip-like isoform [1]. X-ray diffraction data was collected at 1.55 Å resolution (Table 2). The structure contains three GluA2-LBD molecules (chains A-C) in the asymmetric unit of the crystal. Chains A and B form a dimer (Figure 7A), whereas chain C forms a dimer with a symmetry-related chain C.

**Table 2.** X-ray data collection and refinement statistics for GluA2-LBD (L504Y,N775S) with glutamate and compound **32**.

|  |  |
| --- | --- |
| **Crystal data** |  |
| Space group | *P*2<sub>1</sub>2<sub>1</sub>2 |
| Unit cell axes *a*, *b*, *c* (Å) | 114.32, 163.42, 47.47 |
| Molecules in a.u.<sup>a</sup> | 3 |
| **Data collection** |  |
| Beamline | ESRF, ID29 |
| Wavelength (Å) | 0.9763 |
| Resolution (Å) | 49.17-1.55 (1.63-1.55)<sup>b</sup> |
| No. of unique reflections | 129,920 (18,786) |
| Average redundancy | 6.7 (6.6) |
| Completeness (%) | 100.0 (100.0) |
| *R*<sub>merge</sub> (%) | 7.1 (87.9) |
| Mean *I*/σ(*I*) | 13.1 (2.0) |
| Wilson B (Å<sup>2</sup>) | 18.1 |
| **Refinement** |  |
| Amino-acid residues (chain *A/B/C*) | 264/264/264 |
| Glutamate/**32** | 3/3 |
| Water/glycerol/chloride/cacodylate<br>/zinc/acetate | 790/4/3/1<br>/7/8 |
| *R*<sub>work</sub>/*R*<sub>free</sub><sup>c</sup> (%) | 16.8/19.5 |
| *Average B values (Å<sup>2</sup>) for:* |  |
| Amino-acid residues (chain *A/B/C*) | 33.0/29.4/26.1 |
| Glutamate/**32** | 19.0/20.4 |
| Water/glycerol/chloride/cacodylate<br>/zinc/acetate | 34.0/48.8/34.0/25.2<br>/40.0/42.5 |
| RMSD bond length (Å)/angles (°) | 0.007/0.89 |
| Ramachandran outliers/favored (%)<sup>d</sup> | 0.0/98.9 |
| Rotamer outliers (%) /Cβ outliers (%) /Clash score<sup>d</sup> | 1.59/0/1.00 |

<sup>a</sup> A.u.: asymmetric unit of the crystal.
<sup>b</sup> Values in parentheses correspond to the outermost resolution shell.
<sup>c</sup> *R*<sub>free</sub> is equivalent to *R*<sub>work</sub>, but calculated with 5% of reflections omitted from the refinement process.
<sup>d</sup> MolProbity statistics from Phenix.

**Figure 7.**
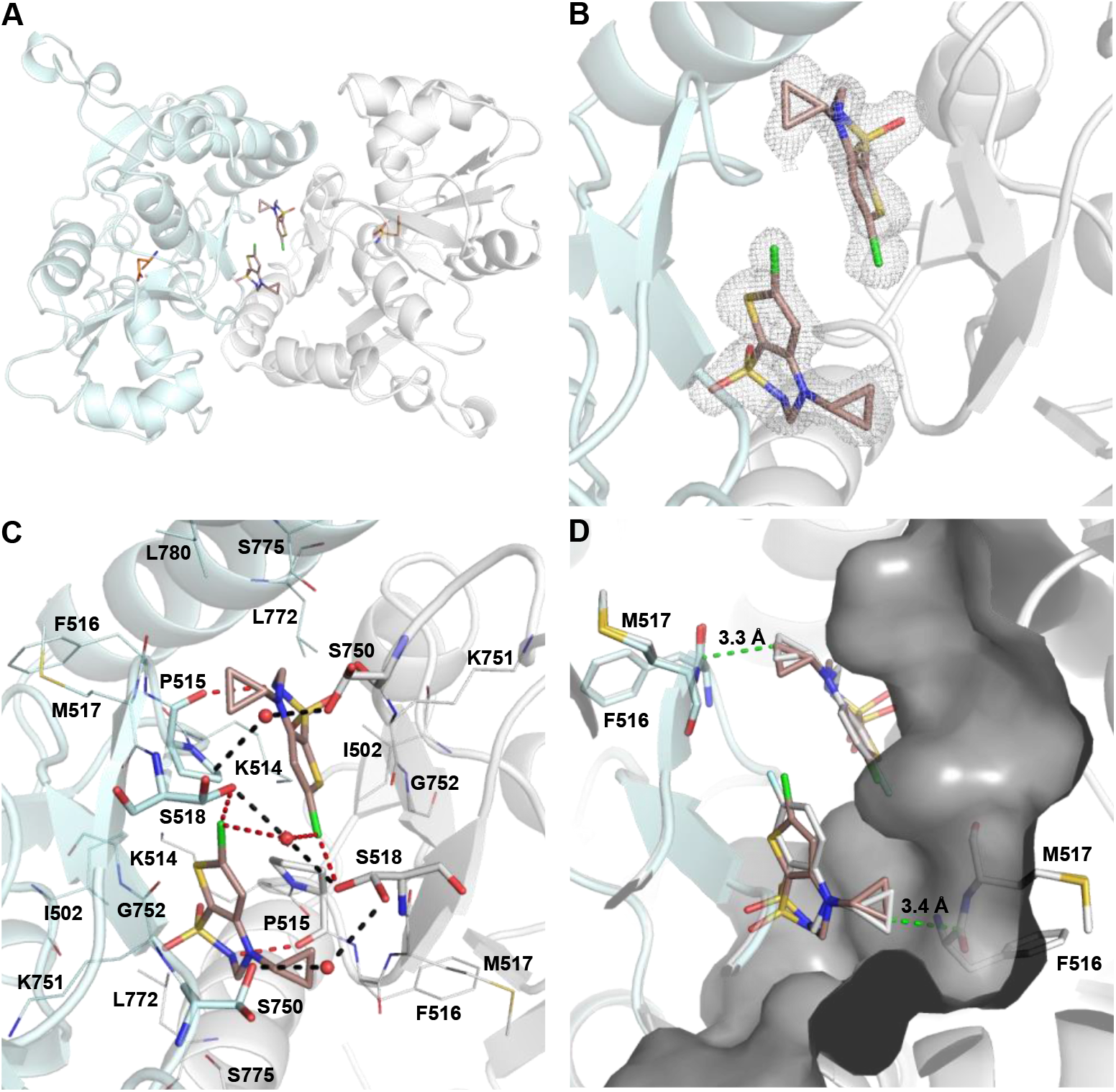
Structure of the ligand-binding domain of GluA2 (L504Y,N775S) in complex with glutamate and **32**. (**A**) Bottom view of the dimeric GluA2-LBD structure (chain A in grey and chain B in palecyan). Glutamate and **32** are shown in sticks representation with carbon atoms in orange and darksalmon, respectively. Oxygen atoms are shown in red, nitrogen atoms in blue, sulfur atoms in yellow, and chlorine atoms in green (**B**) Close-up view on the allosteric binding site with **32**, showing a simple 2Fo-Fc omit electron density map displayed at 1 sigma and carved at 1.6 Å around **32**. (**C**) Close-up view on the allosteric binding site with **32**, showing hydrogen bonding interactions involving **32** (red dashed lines). Residues within 4 Å of the two **32** molecules are shown in lines representation. Pro515, Ser518, and Ser750 in each subunit are shown in sticks representation. The three water molecules taking part in a water-mediated contacts between the two subunits are shown as red spheres. Hydrogen bonds not involving **32** are shown as black dashed lines. (**D**) Comparison of the binding mode of **32** and **10** (PDB code 4N07, chains B and C, protein residues not shown for clarity) in GluA2-LBD. The two structures were aligned on lobe D1 residues (**32**: chain A; **10**: chain B). **10** is shown in sticks representation with carbon atoms in white. Surface for chain A is shown in grey and the shortest distance between the cyclopropyl group of **32** and GluA2 residues are indicated by green dashed lines.

Glutamate is observed in the orthosteric binding site and forms contacts to GluA2 residues as previously observed [47]. Two molecules of **32** bind at an allosteric site in the dimer interface (Figure 7A-C). The electron density for **32** is well-defined and allows unambiguous placement of the modulator (Figure 7B). The binding mode is similar to that of the reference compound **10** [28] (Figure 7D). Binding of **32** does not seem to affect the lobes D1-D2 domain closure (19.0-20.1°) as similar domain closure is observed with glutamate alone [47]. This is in agreement with what has previously been observed for other PAMs [2].

Compound **32** forms a hydrogen bond with the backbone oxygen atom of Pro515 via its sulphonamide nitrogen atom (Figure 7C) as seen in other structures with BTD-type AMPAR PAMs [2]. In addition, the chlorine atom in **32** forms a hydrogen bond with the side chain hydroxyl group of Ser518 from the other subunit (2.9-3.0 Å) and to a water molecule. Ser518 is observed in two conformations in all three GluA2 chains. In the other conformation, the hydroxyl group of Ser518 forms a water-mediated hydrogen bond with the hydroxyl group of Ser750, also observed to adopt two different conformations as well as a water-mediated hydrogen bond to the hydroxyl group of Ser518 in the other subunit. In the more studied compound **10**, a hydrogen bond is formed between a fluorine atom in **10** and Ser518 [28]. It has previously been reported that interaction energies of aliphatic Cl and F atoms are very similar to each other, although the former was slightly higher [48].

The thiophene ring of **32** is located within 4 Å distance of the backbone atoms of Ser750, Lys751, and Gly752 of one subunit and Pro515 of both subunits. The cyclopropyl group of **32** is located within 4 Å of Ser750 as well as Pro515, Phe516, Met517, Ser518, and Leu780 of the other subunit. The sulphonamide oxygen atoms, located within 4 Å of Ile502 and Lys751 as well as Lys514, Pro515, and Leu772 of the other subunit, do not form hydrogen bonds to GluA2 residues or water molecules.

Introduction of a longer substituent at N4 compared to the cyclopropyl group in **32** leads to a decrease in the potency (**27**: EC2x = 19 µM; EC_50_ = 26.9 µM vs. **32**: EC2x = 0.25 µM; EC_50_ = 4.7 µM). A good shape complementarity is observed between the cyclopropyl group and GluA2-LBD, with a closest distance of 3.3 Å (Figure 7D). Therefore, the substituent in **27** would need to adopt a bent conformation, or the modulator would have a different binding mode than **32**. Likewise, cyclization at C3-N4 leads to a decrease in potency (**36**: EC2x = 14 µM; EC_50_ = 40.2 µM vs. **32**: EC2x = 0.25 µM; EC_50_ = 4.7 µM).

Introduction of a methyl group at the sulphonamide N2 leads to a further decrease in the potency (**28**: EC2x = 80 µM; EC_50_ = 76.5 µM vs. **27**: EC2x = 19 µM; EC_50_ = 26.9 µM). In compound **32**, a hydrogen bond is formed between the N2 and Pro515, which would be lost in modulators with a substituent on this nitrogen atom. Introduction of an additional methyl group at C3, besides a methyl group in the N2 position and 2,2-fluoro group in the N4 position, seems to lead to an increase in potency (**24**: EC2x = 1.3 µM; EC_50_ = 31.5 µM vs. **22**: EC2x = 25 µM; EC_50_ = 39.9 µM). This is in line with a previous study reporting that a methyl substituent in the C3 position directs the binding mode of 1,2,4-benzothiadiazine 1,1-dioxide modulators [31].

## 3. Conclusions

We have synthesized new examples of 3,4-dihydro-2*H*-1,2,4-thieno[3,2-*e*]-1,2,4-thiadiazine 1,1-dioxides and examined their potentiator activity on AMPARs and KARs. Particular attention was paid to the introduction of new kinds of alkyl groups at the 4-position (cyclopropyl, allyl) and the preparation of 2,4-dialkyl-/2,3,4-trialkyl-substituted compounds.

The introduction of a methyl group at the 2,3-positions of the heterocycle provided compounds with a marked potentiator activity on AMPARs, while monomethylation at the 2-position appeared to be less favorable (i.e., compare **22** and **24**).

The introduction of a cyclopropyl chain instead of an ethyl chain at the 4-position of the thiadiazine ring was always found to dramatically improve the potentiator activity on AMPARs for previously reported benzo- and pyridothiadiazine dioxides (Figure 2). This structural requirement was clearly confirmed for thienothiadiazine dioxides with compound **32** (BPAM395) expressing a marked *in vitro* activity on AMPARs (EC2x = 0.24 µM) close to that of the 4-cyclopropyl-substituted BTD modulator **10** (BPAM344) used as a reference compound.

Interestingly, compound **27** (BPAM307) emerged as the most promising thienothiadiazine dioxide on KARs being a more efficient potentiator than the 4-cyclopropyl-substituted compound **32** and supporting the view that the allyl side chain at the 4-position of the thiadiazine ring could be more favorable than the cyclopropyl chain to induce marked activity and selectivity for the KARs versus AMPARs.

The tricyclic thieno-analogue **36** of the clinically tested S18986 (**11**) expressed, like the latter, a moderate *in vitro* potentiator activity on AMPARs. However, compound **36** was selected for *in vivo* evaluation in mice as a cognitive enhancer since this compound was found to be safer than **32** after massive *per os* drug administration in mice (100 mg/kg). An increase of the cognition performance of mice was observed with **36** at doses as low as 1 mg/kg *per os*, suggesting that the compound was absorbed after oral administration and reached the CNS to act as a cognitive enhancer.

Finally, the most potent AMPAR potentiator **32** was co-crystallized with the GluA2-LBD (L504Y,N775S) to examine its binding mode at the allosteric binding site located at the LBD dimer interface. It was concluded that this representative of thienothiadiazine dioxide AMPAR PAMs established similar interactions to allosteric binding site residues as the previously reported BTD-type AMPAR modulators.

## 4. Experimental section

### 4.1. Chemistry

#### 4.1.1. General procedures

Melting points were determined on a Büchi Tottoli capillary apparatus and are uncorrected. The ^1^H and ^13^C NMR spectra were recorded on a Bruker Avance (500 MHz for ^1^H; 125 MHz for ^13^C) instrument using deuterated dimethyl sulfoxide (DMSO-d*_6_*) as the solvent with tetramethylsilane (TMS) as an internal standard; chemical shifts are reported in δ values (ppm) relative to that of internal TMS. The abbreviations s = singlet, d = doublet, t = triplet, q = quadruplet, p = pentuplet, m = multiplet, qd = quadruplet of doublet, dt = doublet of triplet, dq = doublet of quadruplet, ddt = doublet of doublet of triplet, td = triplet of doublet, and tt = triplet of triplet are used throughout. Elemental analyses (C, H, N, S) were realized on a Thermo Scientific Flash EA 1112 elemental analyzer and were within ± 0.4% of the theoretical values for carbon, hydrogen, and nitrogen. This analytical method certified a purity of ≥95% for each tested compound. All reactions were routinely checked by TLC on silica gel Merck 60 F254. The synthesis of compounds **14**, **17**, **19**, **20**, **21**, and **25** was previously described [27].

#### 4.1.2. 6-chloro-4-ethyl-2,3-dimethyl-3,4-dihydro-2*H*-thieno[3,2-*e*]-1,2,4-thiadiazine 1,1-dioxide (16)

##### 6-Chloro-4-ethyl-3-methyl-3,4-dihydro-2H-thieno[3,2-e]-1,2,4-thiadiazine 1,1-dioxide (15)

To a solution of 5-chloro-3-(ethylamino)thiophene-2-sulfonamide (**14**) [27] (1.0 g, 4.15 mmol) in acetonitrile (30 mL) was added acetaldehyde (0.7 mL, 0.55 g, 12.5 mmol) and a catalytic amount of camphorsulfonic acid. The reaction mixture was stirred at room temperature for 3-4 h and then evaporated under reduced pressure. The resulting oily residue was triturated with a 1:1 mixture of ethyl acetate and hexane to obtain a white solid, which was collected by filtration and recrystallized in methanol (yields: 30-40%). The unstable compound was immediately used in the next step without further purification.

##### 6-Chloro-4-ethyl-2,3-dimethyl-3,4-dihydro-2H-thieno[3,2-e]-1,2,4-thiadiazine 1,1-dioxide (16)

To the solution of **15** (0.25 g, 0.94 mmol) in acetonitrile (15 mL) was added potassium carbonate (0.5 g, 3.62 mmol) and methyl iodide (0.4 mL, 6.43 mmol). The resulting suspension was heated under stirring at 80°C for 90 min. The reaction mixture was concentrated under reduced pressure and the residue was suspended in a 1:3 mixture of methanol and water. The solid was collected by filtration and recrystallized in methanol (yields: 65-70%). White solid; m.p.: 112-114 °C. ^1^H NMR (DMSO-*d_6_*) δ 1.07 (t, J=7.0 Hz, 3H, NCH_2_C*H_3_*), 1.56 (d, J=6.7 Hz, 3H, CHC*H_3_*), 2.53 (s, 3H, NC*H_3_*), 3.36 (m, 1H, NC*Ha*CH_3_), 3.52 (m, 1H, NC*Hb*CH_3_), 5.18 (q, J=6.7 Hz, 1H, C*H*CH_3_), 7.17 (s, 1H, 5-*H*). ^13^C NMR (DMSO-*d_6_*) δ 13.6 (CH_2_*C*H_3_), 16.9 (CH*C*H_3_), 31.4 (N*C*H_3_), 41.9 (*C*H_2_CH_3_), 71.4 (*C*HCH_3_), 101.0 (C-7a), 117.8 (C-5), 134.5 (C-6), 146.1 (C-4a). Anal. (C_9_H_13_ClN_2_O_2_S_2_) theoretical: C, 38.50; H, 4.67; N, 9.98; S, 22.83. Found: C, 38.40; H, 4.68; N, 10.08; S, 23.14.

#### 4.1.3. 6-Chloro-4-(2,2-difluoroethyl)-2-methyl-3,4-dihydro-2*H*-thieno[3,2-*e*]-1,2,4-thiadiazine 1,1-dioxide (22)

To the solution of 6-chloro-4-(2,2-difluoroethyl)-3,4-dihydro-2*H*-thieno[3,2-*e*]-1,2,4-thiadiazine 1,1-dioxide **21** [27] (0.5 g, 1.73 mmol) in acetonitrile (10 mL) was added potassium carbonate (1.0 g, 7.24 mmol) and methyl iodide (1.0 mL, 16.08 mmol). The resulting suspension was stirred at room temperature for 24 h. The reaction mixture was concentrated under reduced pressure and the residue was suspended in water (30 mL) and extracted with ethyl acetate (3 x 30 mL). The organic layer was washed with water, dried over magnesium sulfate and filtered. The filtrate was concentrated to dryness under reduced pressure. The residue was dissolved in a small volume of ethyl acetate and stirred on an ice bath. The addition of hexane gave rise to the precipitation of the title compound, which was collected by filtration, washed with hexane and dried (yields: 60-65%). White solid; m.p.: 121-123 °C. ^1^H NMR (DMSO-*d_6_*) δ 2.66 (s, 3H, NC*H_3_*), 3.93 (td, J=15.9 Hz/3.5 Hz, 2H, C*H_2_*CHF_2_), 4.95 (s, 2H, NC*H_2_*N), 6.25 (tt, J=55.0 Hz/3.4 Hz, 1H, CH_2_C*H*F_2_), 7.23 (s, 1H, 5-*H*). ^13^C NMR (DMSO-*d_6_*) δ 34.4 (N*C*H_3_), 51.5 (t, J=24 Hz, N*C*H_2_CHF_2_), 67.8 (N*C*H_2_N), 101.4 (C-7a), 114.9 (t, J=241 Hz, NCH_2_*C*HF_2_), 118.1 (C-5), 134.2 (C-6), 146.0 (C-4a). Anal. (C_8_H_9_ClF_2_N_2_O_2_S_2_) theoretical: C, 31.74; H, 3.00; N, 9.25; S, 21.18. Found: C, 31.73; H, 3.06; N, 9.42; S, 21.14.

#### 4.1.4. 6-Chloro-4-(2,2-difluoroethyl)-2,3-dimethyl-3,4-dihydro-2*H*-thieno[3,2-*e*]-1,2,4-thiadiazine 1,1-dioxide (24)

##### 6-Chloro-4-(2,2-difluoroethyl)-3-methyl-3,4-dihydro-2H-thieno[3,2-e]-1,2,4-thiadiazine 1,1-dioxide (23)

To a solution of 5-chloro-3-((2,2-difluoroethyl)amino)thiophene-2-sulfonamide (**19**) [27] (1.0 g, 3.61 mmol) in acetonitrile (20 mL) was added acetaldehyde (1.0 mL, 0.79 g, 17.9 mmol) and a catalytic amount of camphorsulfonic acid. The reaction mixture was heated under stirring at 60 °C for 3-4 h and then evaporated under reduced pressure. The resulting oily residue of the unstable compound **22** was immediately used in the next step without further purification.

##### 6-Chloro-4-(2,2-difluoroethyl)-2,3-dimethyl-3,4-dihydro-2H-thieno[3,2-e]-1,2,4-thiadiazine 1,1-dioxide (24)

To the solution of 6-chloro-4-(2,2-difluoroethyl)-3-methyl-3,4-dihydro-2*H*-thieno[3,2-*e*]-1,2,4-thiadiazine 1,1-dioxide **23** from the previous step in acetonitrile (15 mL) was added potassium carbonate (1.0 g, 7.24 mmol) and methyl iodide (1.0 mL, 16.08 mmol). The resulting suspension was stirred at room temperature for 24 h. The reaction mixture was concentrated under reduced pressure and the residue was extracted with chloroform (50 mL). The organic layer was washed with water and dried over anhydrous magnesium sulfate. The filtrate was concentrated to dryness and the residue of the title compound was recrystallized in methanol (yields of steps 1 and 2: 40-45%). White solid; m.p.: 123-124 °C. ^1^H NMR (DMSO-*d_6_*) δ 1.58 (d, J=6.7 Hz, 3H, CHC*H_3_*), 2.58 (s, 3H, NC*H_3_*), 3.86 (qd, J=15.6 Hz/3.7 Hz, 1H, C*Ha*CHF_2_), 3.99 (qd, J=15.7 Hz/3.6 Hz, 1H, C*Hb*CHF_2_), 5.25 (q, J=6.7 Hz, 1H, C*H*CH_3_), 6.24 (tt, J=54.9 Hz/3.6 Hz, 1H, CH_2_C*H*F_2_), 7.23 (s, 1H, 5-*H*). ^13^C NMR (DMSO-*d_6_*) δ 16.8 (CH*C*H_3_), 31.6 (N*C*H_3_), 48.7 (t, J=24 Hz, N*C*H_2_CHF_2_), 72.6 (*C*HCH_3_), 102.9 (C-7a), 114.9 (t, J=242 Hz, NCH_2_*C*HF_2_), 118.6 (C-5), 134.0 (C-6), 146.1 (C-4a). Anal. (C_9_H_11_ClF_2_N_2_O_2_S_2_) theoretical: C, 34.13; H, 3.50; N, 8.84; S, 20.24. Found: C, 34.11; H, 3.40; N, 9.17; S, 20.47.

#### 4.1.5. 4-Allyl-6-chloro-3,4-dihydro-2*H*-thieno[3,2-*e*]-1,2,4-thiadiazine 1,1-dioxide (27)

##### 4-Allyl-6-chloro-4H-thieno[3,2-e]-1,2,4-thiadiazine 1,1-dioxide (26)

To the solution of 6-chloro-4*H*-thieno[3,2-*e*]-1,2,4-thiadiazine 1,1-dioxide (**25**) [27] (0.68 g, 3.05 mmol) in acetonitrile (30 mL) was added potassium carbonate (1.3 g, 9.41 mmol) and allyl bromide (1.0 mL, 1.4 g, 11.57 mmol). The suspension was heated under stirring for 90 min and then concentrated to dryness under reduced pressure. The residue was taken up in water and the insoluble material was collected by filtration and washed with water. The solid was recrystallized in methanol (yields: 85-90 %). ^1^H NMR (DMSO-*d_6_*) δ 4.67 (dt, J=5.5 Hz/1.6 Hz, 2H, CH_2_CHC*H_2_*), 5.28 (m, 2H, C*H_2_*CHCH_2_), 5.98 (ddt, J=17.3 Hz/10.6 Hz/5.3 Hz, 1H, CH_2_C*H*CH_2_), 7.51 (s, 1H, 5-*H*), 8.10 (s, 1H, 3-*H*).

##### 4-Allyl-6-chloro-3,4-dihydro-2H-thieno[3,2-e]-1,2,4-thiadiazine 1,1-dioxide (27)

The suspension of 4-allyl-6-chloro-4*H*-thieno[3,2-*e*]-1,2,4-thiadiazine 1,1-dioxide (**26**) (0.67 g, 2.55 mmol) in isopropanol (35 mL) was supplemented with sodium borohydride (0.6 g, 15.86 mmol) and the reaction mixture was heated under stirring at 60 °C for 10 min. The solvent was removed by distillation under reduced pressure. The residue was taken up in water and the aqueous suspension was neutralized by 6N HCl and then extracted with chloroform (3 x 30 mL). The combined organic layers were dried over anhydrous magnesium sulfate and filtered. The filtrate was concentrated under reduced pressure and the residue was crystallized in a 1:2 mixture of methanol and water (yields: 80-85 %). White solid; m.p.: 85-86 °C. ^1^H NMR (DMSO-*d_6_*) δ 3.99 (dt, J=5.6 Hz/1.6 Hz, 2H, CH_2_CHC*H_2_*), 4.66 (s, 2H, NC*H_2_*N), 5.18 (dq, J=10.4 Hz/1.5 Hz, 1H, C*Ha*CHCH_2_), 5.26 (dq, J=17.1 Hz/1.7 Hz, 1H, C*Hb*CHCH_2_), 5.84 (ddt, J=17.2 Hz/10.6 Hz/5.5 Hz, 1H, CH_2_C*H*CH_2_), 7.07 (s, 1H, 5-*H*), 8.06 (s, 1H, N*H*). ^13^C NMR (DMSO-*d_6_*) δ 52.4 (*C*H_2_CHCH_2_), 60.9 (N*C*H_2_N), 104.4 (C-7a), 117.4 (CH_2_CH*C*H_2_), 118.2 (C-5), 133.4 (CH_2_*C*HCH_2_), 133.5 (C-6), 147.3 (C-4a). Anal. (C_8_H_9_ClN_2_O_2_S_2_) theoretical: C, 36.29; H, 3.43; N, 10.58; S, 24.22. Found: C, 36.24; H, 3.46; N, 10.44; S, 24.10.

#### 4.1.6. 4-Allyl-6-chloro-2-methyl-3,4-dihydro-2*H*-thieno[3,2-*e*]-1,2,4-thiadiazine 1,1-dioxide (28)

To the solution of 4-allyl-6-chloro-3,4-dihydro-2*H*-thieno[3,2-*e*]-1,2,4-thiadiazine 1,1-dioxide (**27**) (0.3 g, 1.13 mmol) in acetonitrile (10 mL) was added potassium carbonate (0.6 g, 4.34 mmol) and methyl iodide (0.3 mL, 4.82 mmol). The resulting suspension was heated under stirring at 60 °C for 2 h. The reaction mixture was concentrated under reduced pressure and the residue was taken up in water. The insoluble material was collected by filtration, washed with water, and recrystallized in methanol (yields: 60-65%). White solid; m.p.: 90-92 °C. ^1^H NMR (DMSO-*d_6_*) δ 2.64 (s, 3H, NC*H_3_*), 4.04 (dt, J=5.9 Hz/1.6 Hz, 2H, CH_2_CHC*H_2_*), 4.86 (s, 2H, NC*H_2_*N), 5.20 (dq, J=10.3 Hz/1.6 Hz, 1H, C*Ha*CHCH_2_), 5.26 (dq, J=17.3 Hz/1.7 Hz, 1H, C*Hb*CHCH_2_), 5.85 (ddt, J=17.2 Hz/10.2 Hz/5.7 Hz, 1H, CH_2_C*H*CH_2_), 7.12 (s, 1H, 5-*H*). ^13^C NMR (DMSO-*d_6_*) δ 34.8 (N*C*H_3_), 52.6 (*C*H_2_CHCH_2_), 67.0 (N*C*H_2_N), 100.3 (C-7a), 117.7 (CH_2_CH*C*H_2_), 117.7 (C-5), 133.3 (CH_2_*C*HCH_2_), 134.4 (C-6), 146.2 (C-4a). Anal. (C_9_H_11_ClN_2_O_2_S_2_) theoretical: C, 38.78; H, 3.98; N, 10.05; S, 23.00. Found: C, 38.79; H, 3.97; N, 10.18; S, 23.07.

#### 4.1.7. 6-Chloro-4-cyclopropyl-3,4-dihydro-2*H*-thieno[3,2-*e*]-1,2,4-thiadiazine 1,1-dioxide (32)

##### 5-Chloro-3-((1-methoxycyclopropyl)amino)thiophene-2-sulfonamide (29)

To the solution of 3-amino-5-chlorothiophene-2-sulfonamide (**17**) [27] (1.0 g, 4.01 mmol) in methanol (25 mL) was added (1-ethoxycyclopropoxy)trimethylsilane (4 mL, 3.47 g, 19.9 mmol) and glacial acetic acid (4 mL). The reaction mixture was refluxed for 24 h and then introduced in a separating funnel containing ethyl acetate (50 mL). The organic layer was washed with water and dried over anhydrous magnesium sulfate. After filtration, the filtrate was evaporated to dryness under reduced pressure. The residue was taken up with a small volume ethyl acetate and the mixture was cooled at -50 °C. The resulting precipitate was collected by filtration, washed with cooled ethyl acetate, and dried (yields: 35-40 %). ^1^H NMR (DMSO-*d_6_*) δ 0.89 (m, 2H, C(C*H_2_*)_2_), 1.07 (m, 2H, C(C*H_2_*)_2_), 3.19 (s, 3H, OC*H_3_*), 6.91 (s, 1H, N*H*), 6.97 (s, 1H, 4-*H*), 7.55 (s, 2H, SO_2_N*H_2_*).

##### 5-Chloro-3-(cyclopropylamino)thiophene-2-sulfonamide (30)

The solution of 5-chloro-3-((1-methoxycyclopropyl)amino)thiophene-2-sulfonamide (**29**) (0.4 g, 1.41 mmol) in THF (15 mL) was supplemented with sodium borohydride (0.8 g, 21.15 mmol) and boron trifluoride diethyl etherate (0.8 mL). The reaction mixture was refluxed for 12 h and then concentrated to dryness under reduced pressure. The residue was taken up in water and the aqueous suspension was neutralized with 6N HCl. The suspension was extracted with chloroform (3 x 30 mL) and the combined organic layers were dried over anhydrous magnesium sulfate. After filtration, the filtrate was concentrated to dryness under reduced pressure and the residue was dissolved in the minimum ethyl acetate. The addition of hexane provided the formation of a precipitate, which was collected by filtration, washed with hexane, and dried (yields: 75-80 %). ^1^H NMR (DMSO-*d_6_*) δ 0.50 (m, 2H, CH(C*H_2_*)_2_), 0.74 (m, 2H, CH(C*H_2_*)_2_), 2.55 (m, 1H, C*H*(CH_2_)_2_), 5.99 (s, 1H, N*H*), 7.03 (s, 1H, 4-*H*), 7.41 (s, 2H, SO_2_N*H_2_*).

##### 6-Chloro-4-cyclopropyl-4H-thieno[3,2-e]-1,2,4-thiadiazine 1,1-dioxide (31)

The suspension of 5-chloro-3-(cyclopropylamino)thiophene-2-sulfonamide (**30**) (0.27 g, 1.07 mmol) in triethyl orthoformate (7 mL) was heated under stirring at 150 °C in an open vessel for 3-4 h. The reaction mixture was cooled on an ice bath and the precipitate of the title compound was collected by filtration and washed with diethyl ether (yields: 60-65 %). ^1^H NMR (DMSO-*d_6_*) δ 1.07 (m, 4H, CH(C*H_2_*)_2_), 3.40 (m, 1H, C*H*(CH_2_)_2_), 7.59 (s, 1H, 5-*H*), 8.07 (s, 1H, N*H*).

##### 6-Chloro-4-cyclopropyl-3,4-dihydro-2H-thieno[3,2-e]-1,2,4-thiadiazine 1,1-dioxide (32)

The suspension of 6-chloro-4-cyclopropyl-4*H*-thieno[3,2-*e*]-1,2,4-thiadiazine 1,1-dioxide (**31**) (0.17 g, 0.65 mmol) in isopropanol (10 mL) was supplemented with sodium borohydride (0.17 g, 4.49 mmol) and the reaction mixture was heated under stirring at 60 °C for 10 min. The solvent was removed by distillation under reduced pressure. The residue was taken up in water and the aqueous suspension was neutralized by 6N HCl and then extracted with chloroform (3 x 30 mL). The combined organic layers were dried aver anhydrous magnesium sulfate and filtered. The filtrate was concentrated under reduced pressure and the residue was crystallized in a 1:2 mixture of methanol and water (yields: 80-85 %). White solid; m.p.: 168-170 °C. ^1^H NMR (DMSO-*d_6_*) δ 0.68 (m, 2H, CH(C*H_2_*)_2_), 0.82 (m, 2H, CH(C*H_2_*)_2_), 2.63 (m, 1H, C*H*(CH_2_)_2_), 4.63 (s, 2H, NC*H_2_*N), 7.13 (s, 1H, 5-*H*), 7.99 (s, 1H, N*H*). ^13^C NMR (DMSO-*d_6_*) δ 7.6 (CH(*C*H_2_)_2_), 31.1 (*C*H(CH_2_)_2_), 61.7 (N*C*H_2_N), 105.7 (C-7a), 118.2 (C-5), 133.4 (C-6), 148.4 (C-4a). Anal. (C_8_H_9_ClN_2_O_2_S_2_) theoretical: C, 36.29; H, 3.43; N, 10.58; S, 24.22. Found: C, 36.32; H, 3.42; N, 10.72; S, 24.40.

#### 4.1.8. 2-Chloro-5a,6,7,8-tetrahydro-5*H*-pyrrolo[2,1-*c*]thieno[3,2-*e*]-1,2,4-thiadiazine 4,4-dioxide (36)

##### 4-Chloro-N-(5-chloro-2-sulfamoylthiophen-3-yl)butanamide (33)

The solution of 3-amino-5-chlorothiophene-2-sulfonamide (**17**) [27] (4.0 g, 16.04 mmol) and 4-chlorobutyryl chloride (4 mL, 5.03 g, 35.69 mmol) in dioxane (60 mL) was heated under stirring at 70 °C for 90 min. The solvent was removed by distillation under reduced pressure and the oily residue was taken up in water. The resulting suspension was cooled on an ice bath and the insoluble material was collected by filtration, washed with water, and dried (yields: 85-90 %). ^1^H NMR (DMSO-*d_6_*) δ 2.03 (p, J=6.8 Hz, 2H, COCH_2_C*H_2_*CH_2_Cl), 2.57 (t, J=7.4 Hz, 2H, COC*H_2_*CH_2_CH_2_Cl), 3.71 (t, J=6.5 Hz, 2H, COCH_2_CH_2_C*H_2_*Cl), 7.70 (s, 1H, 4-*H*), 7.85 (s, 2H, SO_2_N*H_2_*), 9.29 (s, 1H, N*H*).

##### 5-Chloro-3-(2-oxopyrrolidin-1-yl)thiophene-2-sulfonamide (34)

4-Chloro-*N*-(5-chloro-2-sulfamoylthiophen-3-yl)butanamide (**33**) (4.2 g, 13.24 mmol) was dissolved in 1N NaOH (50 mL) and the solution was stirred at room temperature for 12 h. The reaction mixture was cooled on an ice bath and acidified with 6N HCl. The resulting precipitate of the title compound was collected by filtration, washed with water, and dried (yields: 90-95 %). ^1^H NMR (DMSO-*d_6_*) δ 2.09 (p, J=7.4 Hz, 2H, 4’-*H_2_*), 2.41 (t, J=8.0 Hz, 2H, 3’-*H_2_*), 3.76 (t, J=7.0 Hz, 2H, 5’-*H_2_*), 7.30 (s, 1H, 4-*H*), 7.59 (s, 2H, SO_2_N*H_2_*).

##### 2-Chloro-7,8-dihydro-6H-pyrrolo[2,1-c]thieno[3,2-e]-1,2,4-thiadiazine 4,4-dioxide (35)

5-Chloro-3-(2-oxopyrrolidin-1-yl)thiophene-2-sulfonamide (**34**) (3.3 g, 11.75 mmol) was introduced in an open vessel and heated at 210 °C until the melting of the solid. After 150 min, the reaction mixture was cooled at room temperature and taken up with methanol under reflux. The methanolic solution was cooled on an ice bath and the precipitate of the title compound was collected by filtration, washed with methanol, and dried (yields: 75-80 %). ^1^H NMR (DMSO-*d_6_*) δ 2.18 (p, J=7.6 Hz, 2H, 7-*H_2_*), 2.93 (t, J=7.9 Hz, 2H, 6-*H_2_*), 4.11 (t, J=7.2 Hz, 2H, 8-*H_2_*), 7.48 (s, 1H, 1-*H*).

##### 2-Chloro-5a,6,7,8-tetrahydro-5H-pyrrolo[2,1-c]thieno[3,2-e]-1,2,4-thiadiazine 4,4-dioxide (36)

The suspension of 2-chloro-7,8-dihydro-6*H*-pyrrolo[2,1-*c*]thieno[3,2-*e*]-1,2,4-thiadiazine 4,4-dioxide (**35**) (2.36 g, 8.98 mmol) in isopropanol (40 mL) was supplemented with sodium borohydride (2.0 g, 52.82 mmol) and the reaction mixture was heated under stirring at 60 °C for 5-10 min. The solvent was removed by distillation under reduced pressure. The residue was taken up in water and the aqueous suspension was neutralized by 6N HCl and then extracted with chloroform (3 x 50 mL). The combined organic layers were dried over anhydrous magnesium sulfate and filtered. The filtrate was concentrated under reduced pressure and the residue of the title compound was recrystallized in methanol (yields: 50-55 %). White solid; m.p.: 215-217 °C. ^1^H NMR (DMSO-*d_6_*) δ 0.68 (m, 2H, CH(C*H_2_*)_2_), 0.82 (m, 2H, CH(C*H_2_*)_2_), 2.63 (m, 1H, C*H*(CH_2_)_2_), 4.63 (s, 2H, NC*H_2_*N), 7.13 (s, 1H, 5-*H*), 7.99 (s, 1H, N*H*). ^13^C NMR (DMSO-*d_6_*) δ 7.6 (CH(*C*H_2_)_2_), 31.1 (*C*H(CH_2_)_2_), 61.7 (N*C*H_2_N), 105.7 (C-7a), 118.2 (C-5), 133.4 (C-6), 148.4 (C-4a). Anal. (C_8_H_9_ClN_2_O_2_S_2_) theoretical: C, 36.29; H, 3.43; N, 10.58; S, 24.22. Found: C, 36.28; H, 3.45; N, 10.91; S, 24.25.

#### 4.1.9. 2-Chloro-5-methyl-5a,6,7,8-tetrahydro-5*H*-pyrrolo[2,1-*c*]thieno[3,2-*e*]-1,2,4-thiadiazine 4,4-dioxide (37)

To the solution of 2-chloro-5a,6,7,8-tetrahydro-5*H*-pyrrolo[2,1-*c*]thieno[3,2-*e*]-1,2,4-thiadiazine 4,4-dioxide (**36**) (0.25 g, 0.94 mmol) in acetonitrile (10 mL) was added potassium carbonate (0.5 g, 3.62 mmol) and methyl iodide (0.3 mL, 4.82 mmol). The resulting suspension was heated under stirring at 70 °C for 2-3 h. The reaction mixture was concentrated under reduced pressure and the residue was taken up in water. The insoluble material was collected by filtration, washed with water, and recrystallized in methanol (yields: 80-85%). White solid; m.p.: 147-148 °C. ^1^H NMR (DMSO-*d_6_*) δ 1.96 (m, 1H, 6-*Ha*), 2.06 (m, 1H, 6-*Hb*), 2.15 (m, 1H, 7-*Ha*), 2.32 (m, 1H, 7-*Hb*), 2.36 (s, 3H, C*H_3_*), 3.36 (m, 1H, 8-*Ha*), 3.62 (m, 1H, 8-*Hb*), 5.42 (t, J=7.1 Hz, 5a-*H*), 7.07 (s, 1H, 1-*H*). ^13^C NMR (DMSO-*d_6_*) δ 22.8 (C-7), 26.9 (C-6), 29.0 (*C*H_3_), 48.6 (C-8), 74.6 (C-5a), 98.2 (C-3a), 117.8 (C-1), 134.8 (C-2), 145.1 (C-9a). Anal. (C_9_H_11_ClN_2_O_2_S_2_) theoretical: C, 38.78; H, 3.98; N, 10.05; S, 23.00. Found: C, 38.47; H, 3.92; N, 10.36; S, 22.76.

### 4.2. Biological evaluation

#### 4.2.1. Effect on AMPA-evoked membrane depolarization (*in vitro* fluorescence assay) on rat primary brain cultures

The effect was investigated using an AMPA-evoked membrane depolarization assay, as measured by fluorescent membrane potential dyes and an imaging-based plate reader on rat primary brain cultures, following our previously published procedure [25].

#### 4.2.2. Effect on the calcium influx on HEK293 cells stably expressing the GluA2(*Q*)_i_ subunit

Human Embryonic Kidney Grip Tite^TM^ (HEK293-GT) cells (Invitrogen) was used to create a polyclonal cell line expressing GluA2(*Q*)_i_ as previously described [49]. In short, a pIRES plasmid vector containing cDNA encoding blasticidin-S deaminase enzymes was inserted with the cDNA of rat GluA2(*Q*)_i_. The cell line was maintained in Dulbeccós Modified Eagle Medium (DMEM with GlutaMAX) supplemented with 10% fetal bovine serum, 100 units/mL penicillin, 100 µg/mL streptomycin, 1% geneticin (G418), and 5 µg/mL blasticidin (Sigma-Aldrich) at 37 °C in a humidified 5% CO_2_ incubator. Cells were then cultured in monolayer and grown until 90% confluence prior to use. Cells were seeded into black/clear flat bottom 96-well plates (Greiner, In Vitro) coated with poly-D-lysine and incubated for two days at 37 °C (5% CO_2_) to reach full confluence. Assay plates for detection of intracellular calcium were washed twice in phosphate buffered saline (PBS) before incubated (37 °C for at least 30 min) with DMEM containing 2 µM of FluoAM-8 dye (AAT Bioquest) per well. The tested compounds were dissolved in DMSO and prepared as compound solutions in assay buffer (100 mM choline chloride, 10 mM sodium chloride, 5 mM potassium chloride, 1 mM magnesium chloride, 20 mM calcium chloride, 10 mM HEPES, pH 7.4). Clear V-bottom 96-well plates (Greiner, In Vitro) were used to prepare the compound solutions containing the tested compounds in concentration range 0.2-600 µM in the presence of 100 µM glutamate as agonist. After incubation in dye solution, the cells were washed three times with PBS and 50 µL of assay buffer was added per well prior to detection with a FlexStation I Plate Reader (Molecular Devices). GluA2 receptor activation from application of compound solution was detected as relative fluorescence unit (RFU) at an emission wavelength of 538 nm after excitation at 485 nm. Data was recorded as the peak fluorescence response and subtracted from background fluorescence using the program SoftMax Pro version 5.4 (Molecular Devices). For construction of dose-response curves, the responses from at least three individual experiments with the same modulator concentration (triplicate wells) were normalized to the response at maximal modulator concentration using GraphPad Prism version 9.3.0. Normalized data was used to fit dose-response curves by applying the equation: log(agonist) vs. response – variable slope (four parameters) constraining the top to 100% and the bottom to 0%. The top four modulators with highest potency at GluA2 were also tested at GluK1(*Q*)_1b_ and GluK2(*Q*)_2a_ in the presence of 100 µM KA and 100 µM of the modulator. SigmaPlot version 14.0 was used to make the bar histograms and perform Student’s t-test.

#### 4.2.3. *In vivo* effect on the body temperature in NMRI mice

The effect of the new compounds on the body temperature in male NMRI mice was measured after *per os* administration (30 and 100 mg/kg) according to previously described procedures [50–52].

#### 4.2.4. Effect on long-term potentiation (LTP) of the postsynaptic response evoked in the dentate gyrus on anesthetized rats

Extracellular excitatory postsynaptic field potentials (EPSfP) were recorded in the dentate gyrus using our previously published procedure [25].

#### 4.2.5. Effect in object recognition test in mice

The one-trial object recognition paradigm measures a form of episodic memory in the mouse and was achieved following our previously described procedure [24,25]. The effects of compound **36** was measured in CD1 mice after *per os* administration (1 and 3 mg/kg).

### 4.3. Structure determination

Rat GluA2_o_-LBD (L504Y,N775S) was expressed and purified as previously described [26,53]. A protein-ligand solution containing 4 mg/mL GluA2-LBD, 5 mM glutamate, and compound **32**, added in excess as solid compound to ensure a saturated concentration, was used for crystallization. The protein-ligand buffer contained 10 mM HEPES at pH 7, 20 mM sodium chloride, and 1 mM EDTA. The protein was crystallized using the hanging drop vapor diffusion method. The hanging drop consisted of 1 μL protein-ligand solution plus 1 μL crystallization solution (reservoir). The volume of the reservoir solution was 500 μL. Crystallization of the complex was performed at 6 °C. The crystal used for data collection was obtained using a reservoir solution consisting of 15.2% PEG4000, 0.1 M zinc acetate, 0.1 M sodium cacodylate, pH 6.5. The crystals were cryo-protected using reservoir solution containing 20% glycerol before flash-cooling in liquid nitrogen. X-ray diffraction data was collected at beamline ID29, ESRF, France. The data was processed with XDS [54] and scaled using Scala within the CCP4i programme [55]. The structure was solved with molecular replacement using Phaser in CCP4i and the structure of GluA2-LBD with glutamate and cyclothiazide (PDB code 3TKD, chain A) as search model. The structure was initially refined using Autobuild in Phenix [56]. The program Maestro [Release 2021-3, Schrödinger, LLC, New York, NY, 2021] was used to generate coordinate files for **32**. The modulator parameter file for refinements in Phenix were obtained using eLBOW [57], keeping the geometry obtained from geometry optimization in MacroModel [Maestro Release 2021-3, Schrödinger, LLC, New York, NY, 2021.]. The structure was manually adjusted in Coot [58] and refinement rounds were performed in Phenix with individual isotropic B-factors, TLS, and riding H atoms. GluA2-LBD domain closures were calculated using the DynDom server [59] relative to the apo structure of GluA2_o_-LBD (PDB code 1FTO, chain A). Figure 7 was prepared in PyMOL [version 2.5.5, The PyMOL Molecular Graphics Systems, V.S., Schrödinger LLC].

### 4.4. Accession code

The structure coordinates and corresponding structure factor file of GluA2_o_-LBD (L504Y,N775S) with glutamate and compound **32** have been deposited in the Protein Data Bank (PDB) under the accession code 8QEZ.

**Scheme 1.**
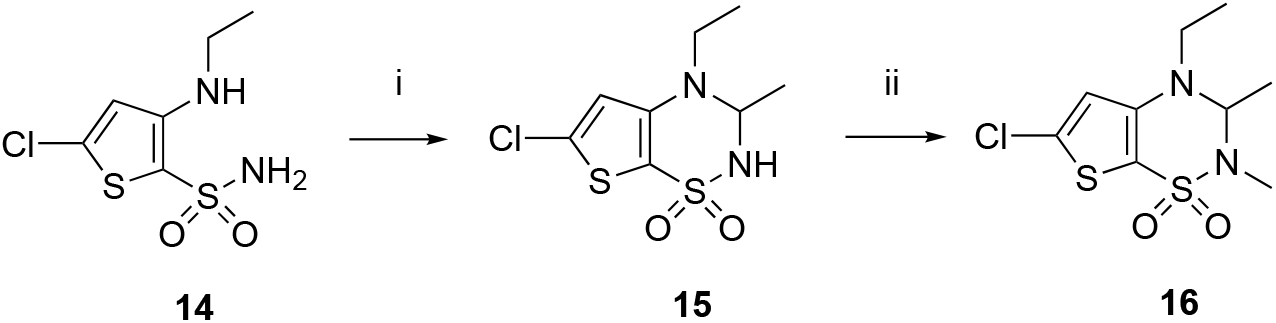
Synthetic pathway to the target compound **16**.

#### Reagents

(i) CH_3_CHO, H^+^, CH_3_CN, r.t., 3-4 h; (ii) CH_3_I, K_2_CO_3_, CH_3_CN, 80 °C, 1−2 h.

**Scheme 2.**
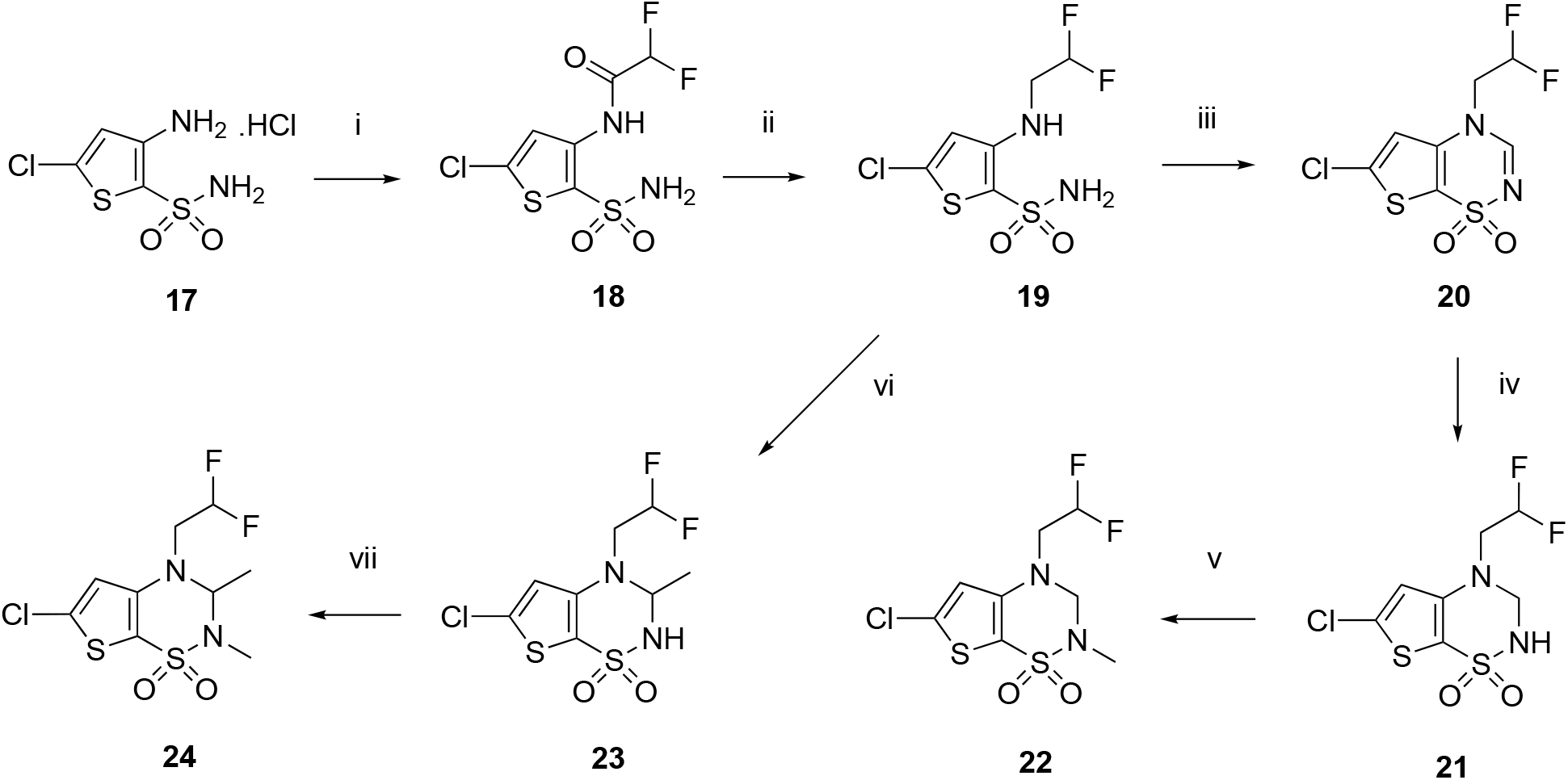
Synthetic pathway to the target compounds **22** and **24**.

#### Reagents

(i) CHF_2_COCl, dioxane, r.t., 72 h; (ii) LiAlH_4_, diethyl ether, 1 h; (iii) HCOOH, 70-80 °C, 3-4 h; (iv) NaBH_4_, isopropanol, 50−55 °C, 30 min; (v) CH_3_I, K_2_CO_3_, CH_3_CN, r.t., 12 h; (vi) CH_3_CHO, H^+^, CH_3_CN, 60 °C, 3-4 h; (vii) CH_3_I, K_2_CO_3_, CH_3_CN, r.t., 12 h.

**Scheme 3.**
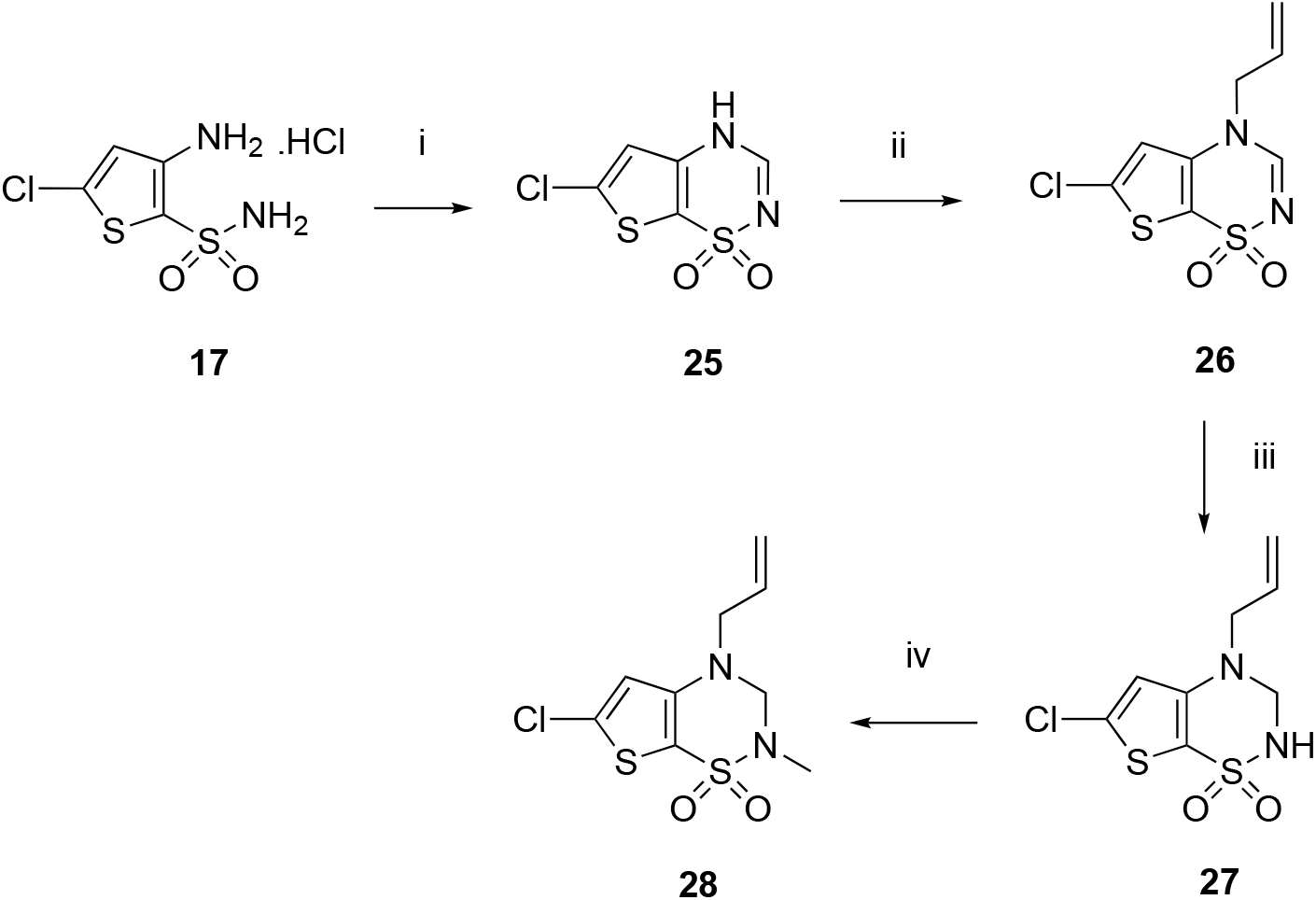
Synthetic pathway to the target compounds **27** and **28**.

#### Reagents

(i) HC(OEt)_3_, 120-130°C, 3 h; (ii) CH_2_=CHCH_2_Br, K_2_CO_3_, CH_3_CN, 65 °C, 90 min; (iii) NaBH_4_, isopropanol, 60 °C, 10 min; (iv) CH_3_I, K_2_CO_3_, CH_3_CN, 80 °C, 2 h.

**Scheme 4.**
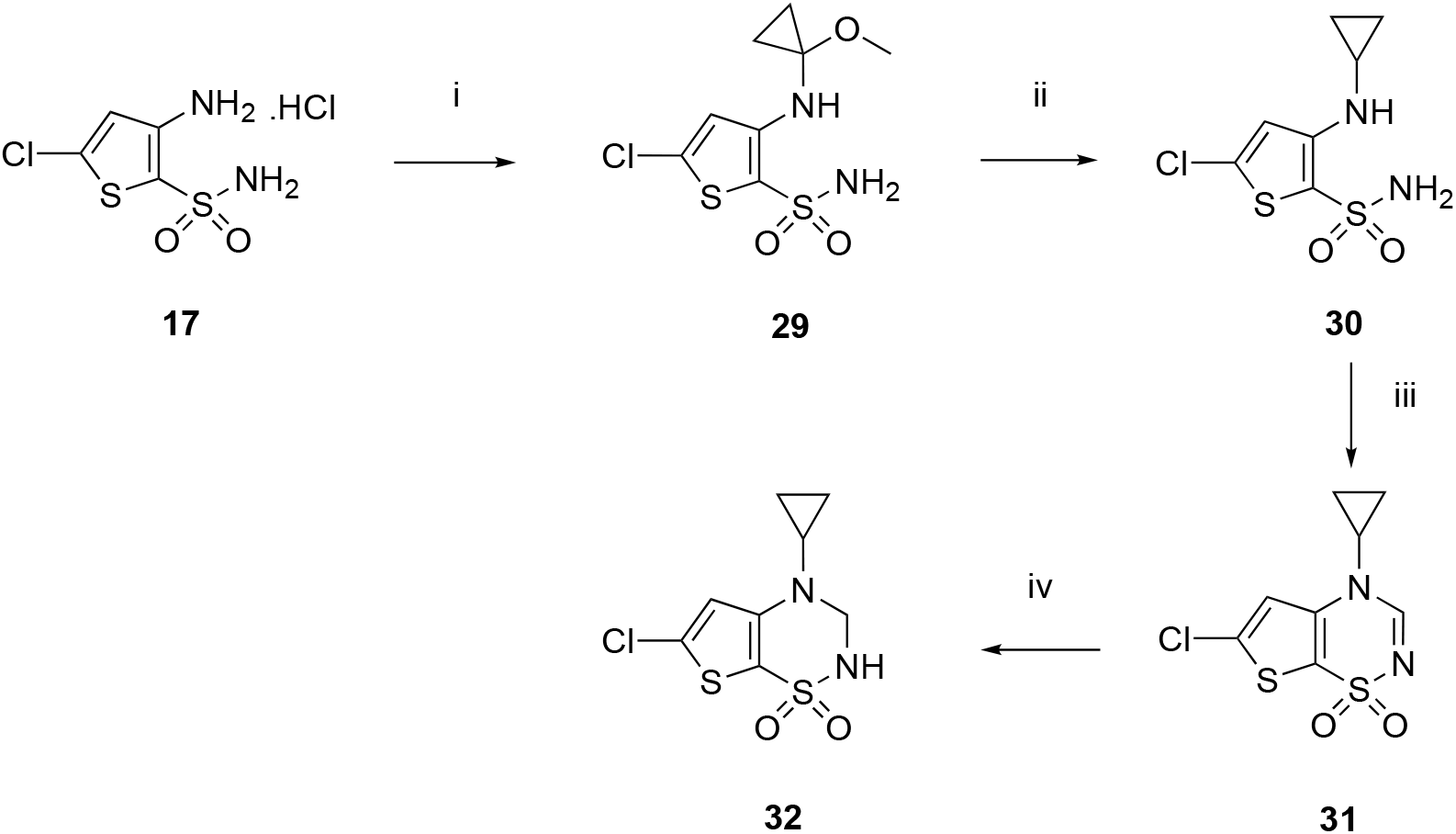
Synthetic pathway to the target compound **32**.

#### Reagents

(i) (1-ethoxycyclopropoxy)trimethylsilane, MeOH, HOAc, reflux, 16 h; (ii) NaBH_4_, BF_3_.Et_2_O, THF, reflux, 12 h; (iii) HC(OEt)_3_, 150°C, 3-4 h; (iv) NaBH_4_, isopropanol, 50−55 °C, 30 min.

**Scheme 5.**
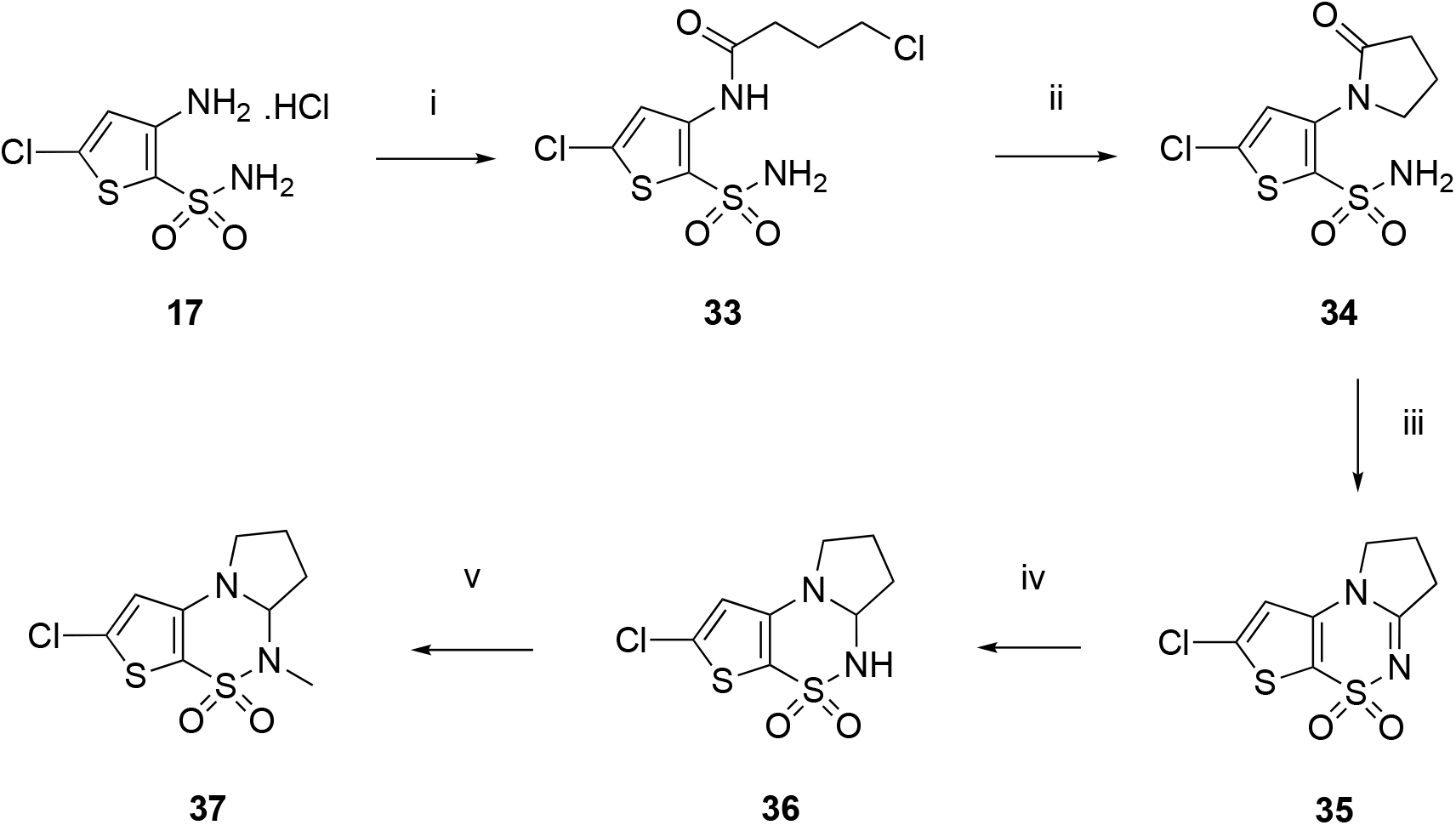
Synthetic pathway to the target compounds **36** and **37**.

#### Reagents

(i) CH_2_ClCH_2_CH_2_COCl, dioxane, 70 °C, 90 min; (ii) NaOH 1N, 12 h; (iii) fusion, 200-210 °C, 2-3 h; (iv) NaBH_4_, isopropanol, 55 °C, 10 min; (v) CH_3_I, K_2_CO_3_, CH_3_CN, 70-80 °C, 2-3 h.

## Supplementary material

Supplementary data to this article (^1^H and ^13^C NMR spectra of the target compounds) can be found online at …

## Acknowledgements

This study was supported in part by a grant from Servier and the National Fund for Scientific Research (F.N.R.S., Belgium) from which P. de Tullio is a Research Director. Y.B., J.D., S.L., and J.S.K. acknowledge funding from the Independent Research Fund Denmark and J.D., S.L., K.F., and J.S.K. from Danscatt.

The technical assistance of S. Counerotte is gratefully acknowledged. We would like to thank Heidi Peterson for expression and purification of GluA2-LBD and beamline scientists at ESRF for their valuable assistance and support in using beamline ID29. We acknowledge the European Synchrotron Radiation Facility (ESRF) for provision of synchrotron radiation facilities under proposal number MX1901.

## Abbreviations

AMPA: α-amino-3-hydroxy-5-methyl-4-isoxazolepropionic acid
AMPAR: AMPA receptor
PAM: positive allosteric modulator
BDNF: brain-derived neurotrophic factor
BTD: 3,4-dihydro-2*H*-1,2,4-benzothiadiazine 1,1-dioxide
CTD: C-terminal domain
EPSfP: excitatory postsynaptic field potential
HEK293-GT: Human Embryonic Kidney Grip Tite^TM^
FlipR: fluorescent imaging plate reader
GluA2-LBD: ligand-binding domain of GluA2 (L504Y,N775S)
iGluR: ionotropic glutamate receptor
KA: kainic acid
KAR: KA receptor
LBD: ligand-binding domain
LTP: long-term potentiation
mGluR: metabotropic glutamate receptor
NMDA: *N*-methyl-*D*-aspartic acid
NTD: N-terminal domain
TMD: transmembrane domain
TMS: tetramethylsilane

