## Supplementary information for "Exploring thienothiadiazine dioxides as isosteric analogues of benzo- and pyridothiadiazine dioxides in the search of new AMPA and kainate receptor positive allosteric modulators"

### Supplementary material

$^{1}$H and $^{13}$C NMR spectra of the target compounds **16**, **22**, **24**, **27**, **28**, **32**, **36**, **37**

$^{1}$H NMR spectra of the intermediates **26**, **29**, **30**, **31**, **33**, **34**, **35**

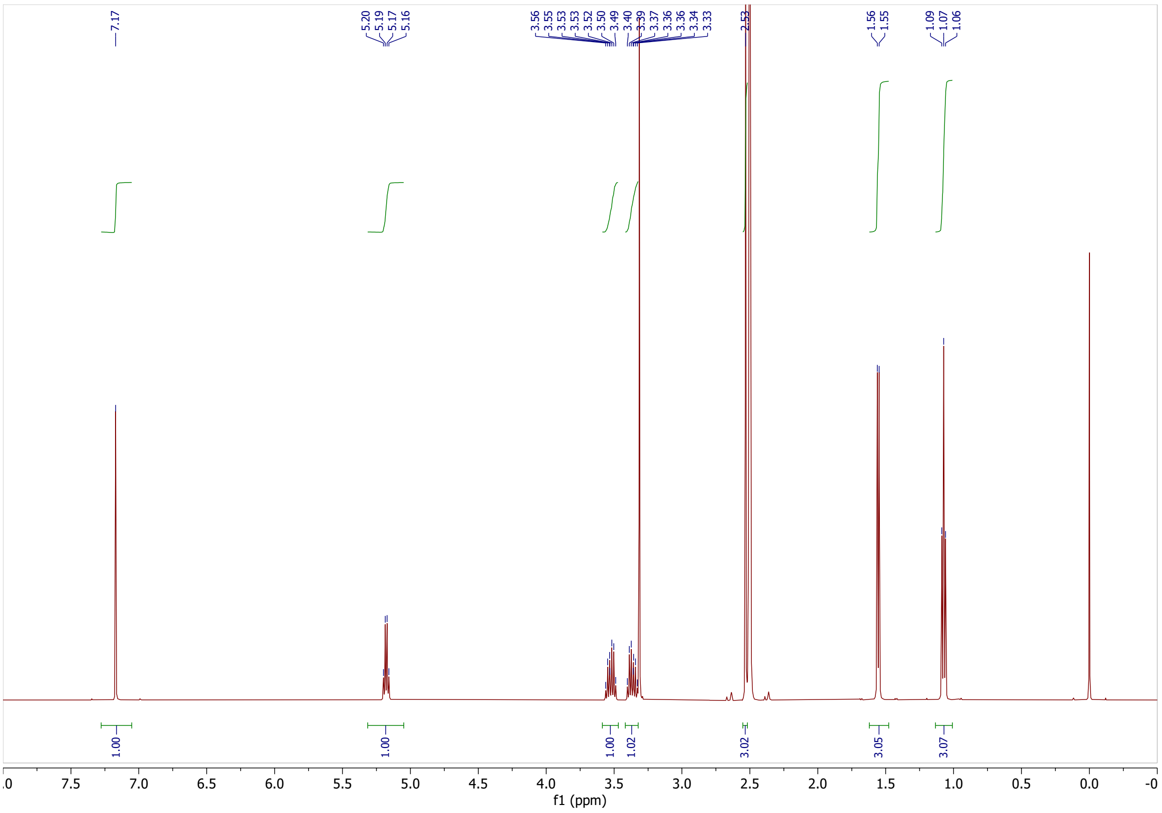

7.17
5.20 5.19 5.17 5.16
3.56 3.55 3.53 3.53 3.52 3.50 3.49 3.40 3.39 3.37 3.36 3.36 3.34 3.33
2.53
1.56 1.55
1.09 1.07 1.06
1.00 1.00 1.00 1.02 3.02 3.05 3.07
f1 (ppm)


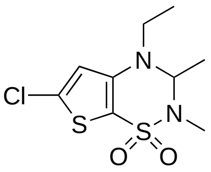

Cl
S
N
N
S
O
O


**16**

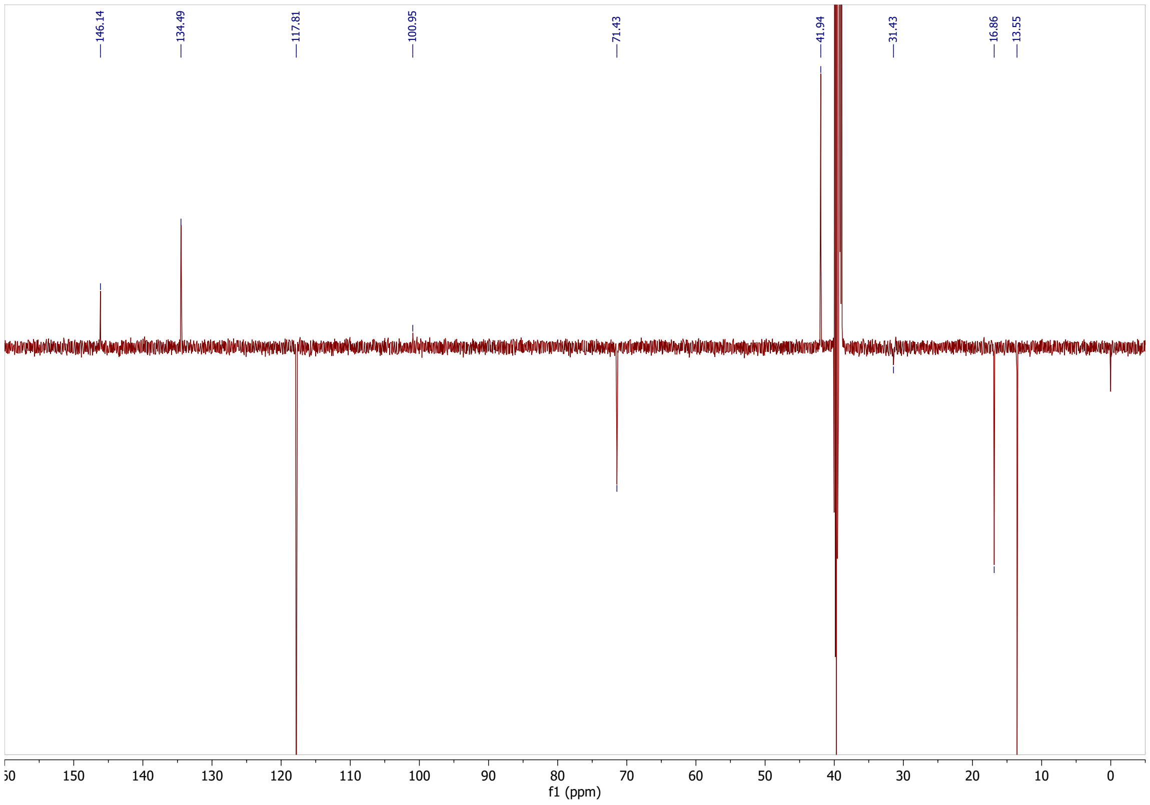

146.14
134.49
117.81
100.95
71.43
41.94
31.43
16.86
13.55
f1 (ppm)

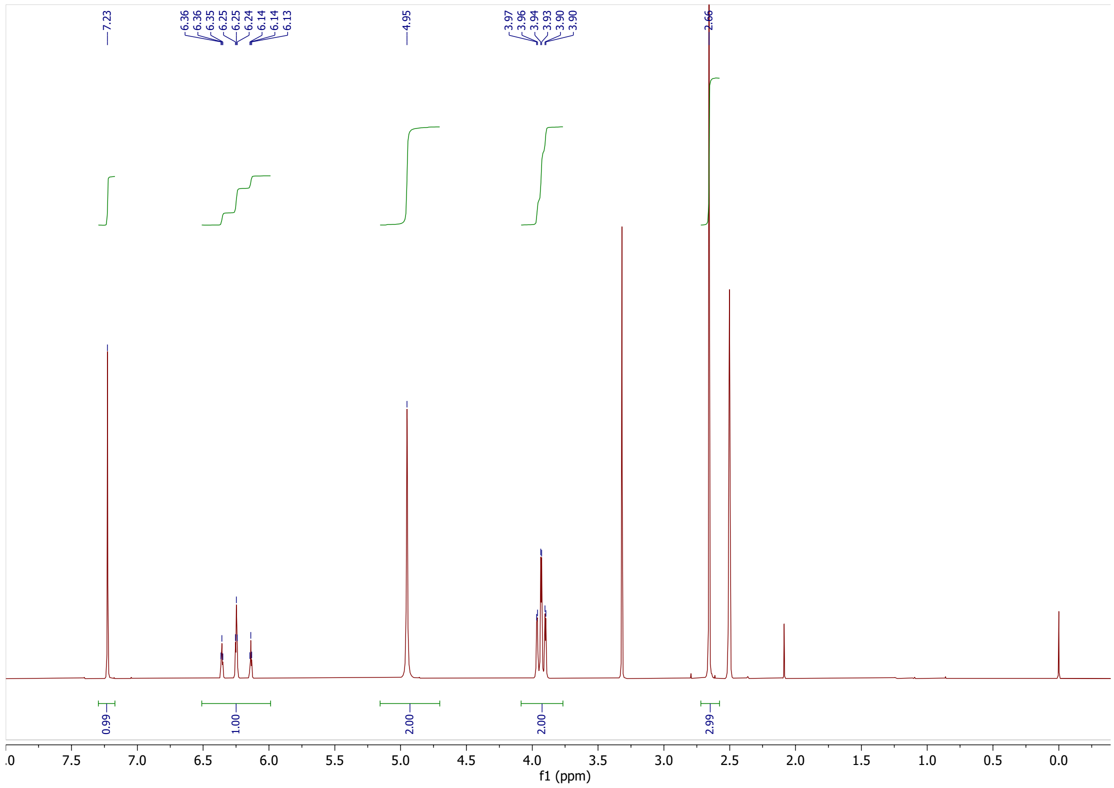
7.23
6.36
6.36
6.35
6.25
6.25
6.24
6.14
6.14
6.13
4.95
3.97
3.96
3.94
3.93
3.90
3.90
2.66
0.99
1.00
2.00
2.00
2.99
7.5
7.0
6.5
6.0
5.5
5.0
4.5
4.0
3.5
3.0
2.5
2.0
1.5
1.0
0.5
0.0
f1 (ppm)

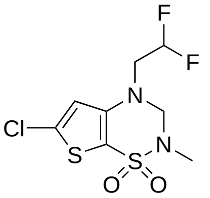
F
F
N
Cl
S
S
N
O
O

**22**

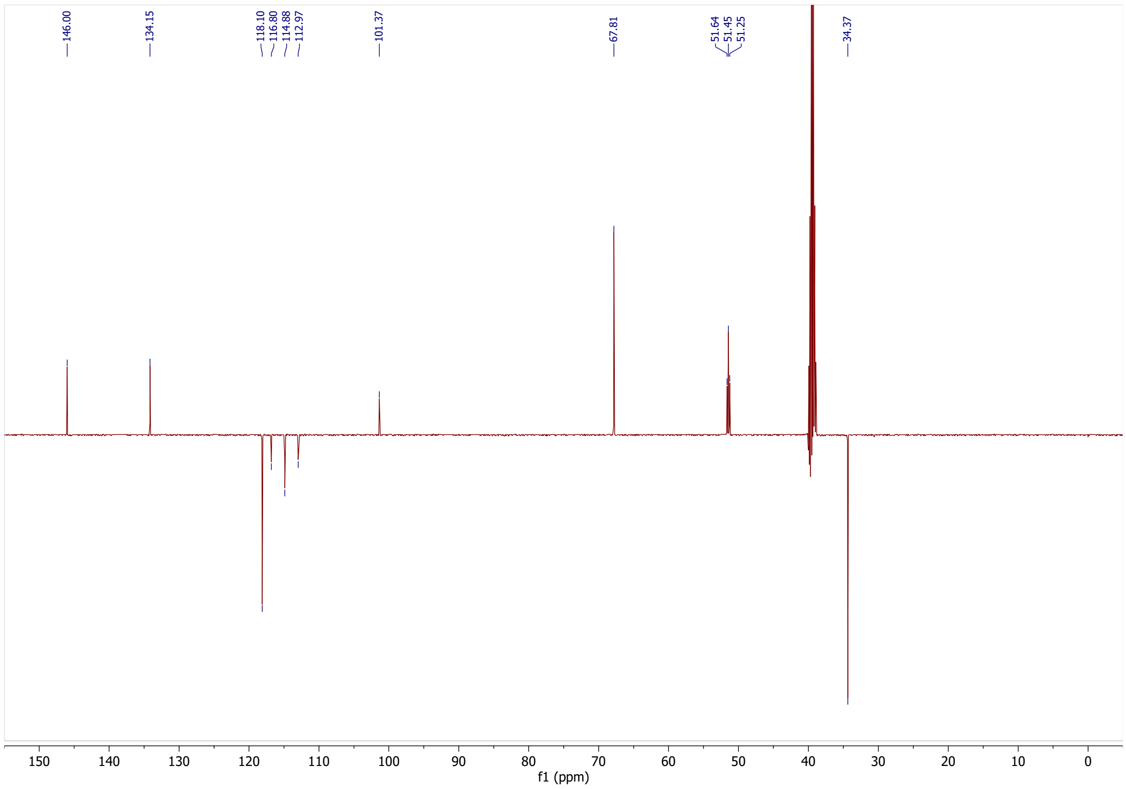
146.00
134.15
118.10
116.80
114.88
112.97
101.37
67.81
51.64
51.45
51.25
34.37
150
140
130
120
110
100
90
80
70
60
50
40
30
20
10
0
f1 (ppm)

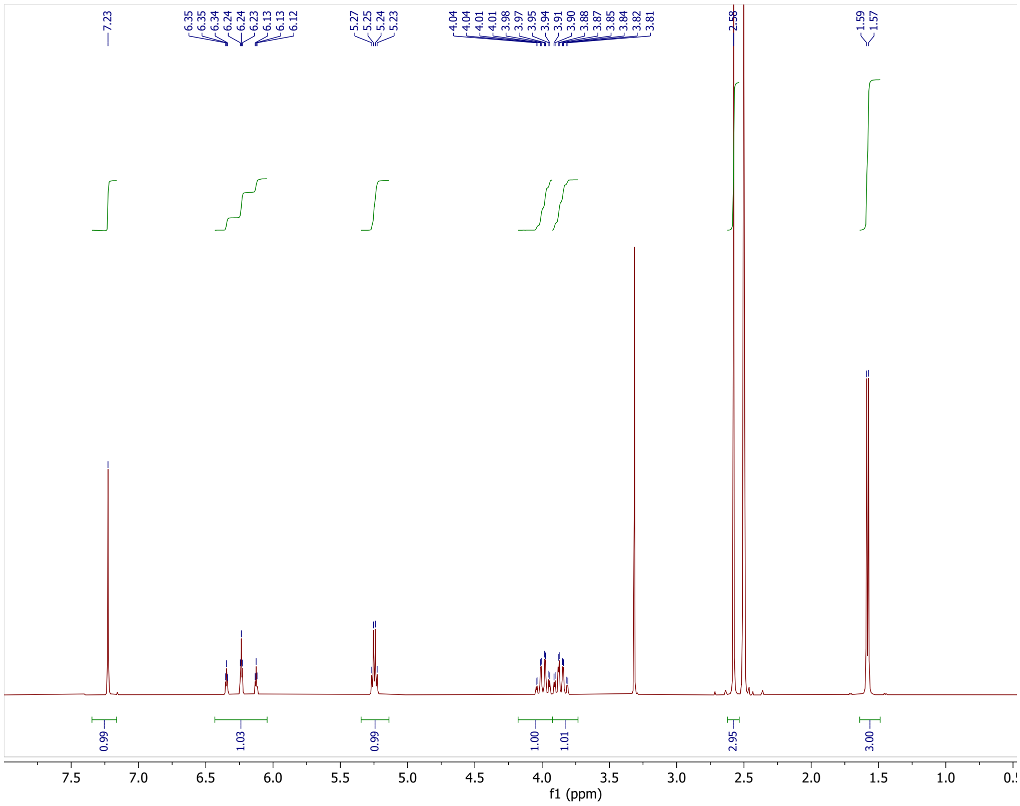
7.23
6.35 6.35 6.34 6.24 6.24 6.23 6.13 6.13 6.12
5.27 5.25 5.24 5.23
4.04 4.04 4.01 4.01 3.98 3.97 3.95 3.94 3.91 3.90 3.88 3.87 3.85 3.84 3.82 3.81
2.58
1.59 1.57
0.99 1.03 0.99 1.00 1.01 2.95 3.00
7.5 7.0 6.5 6.0 5.5 5.0 4.5 4.0 3.5 3.0 2.5 2.0 1.5 1.0
f1 (ppm)

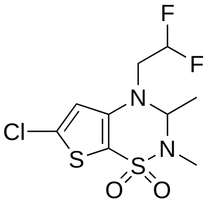
F
F
N
N
Cl
S
S
O
O

**24**

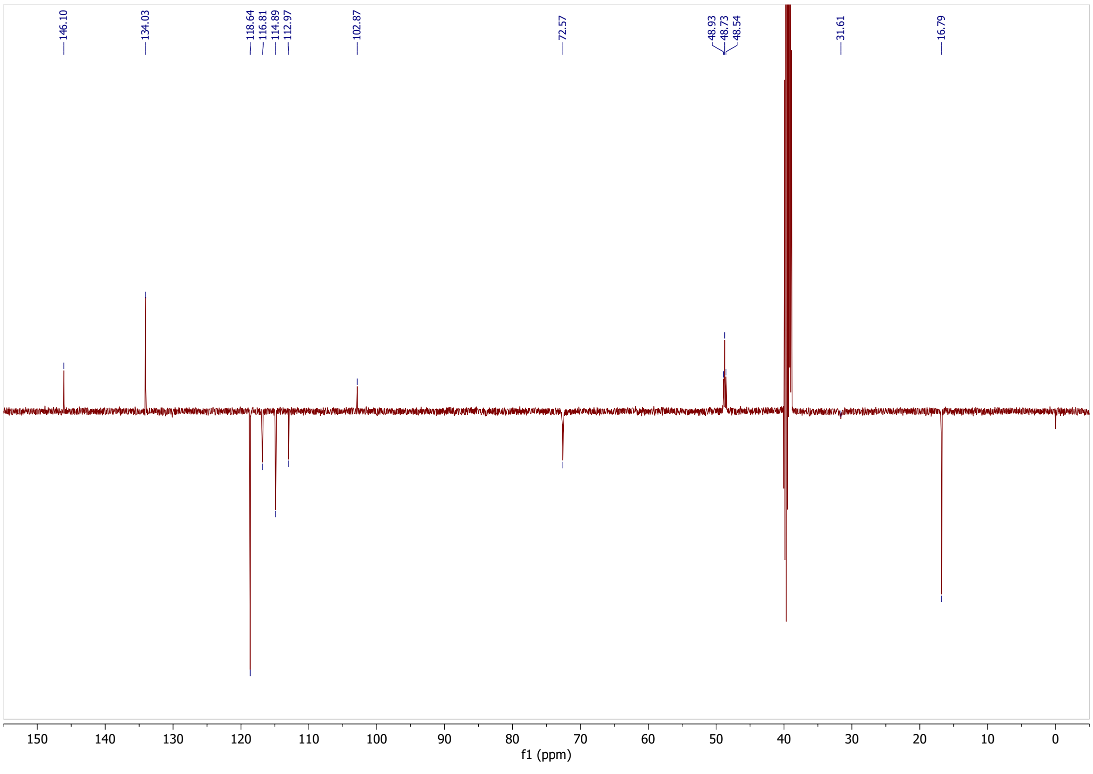
146.10
134.03
118.64 116.81 114.89 112.97
102.87
72.57
48.93 48.73 48.54
31.61
16.79
150 140 130 120 110 100 90 80 70 60 50 40 30 20 10 0
f1 (ppm)

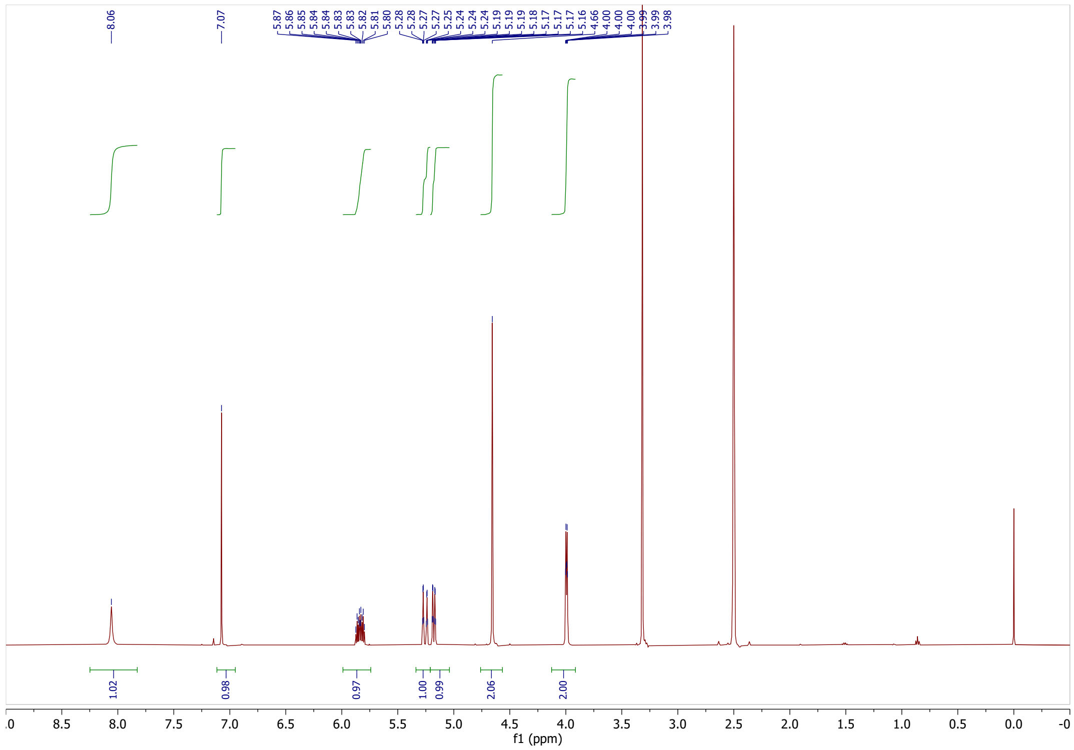

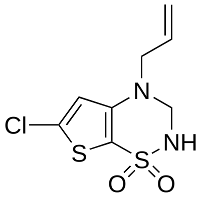


**27**

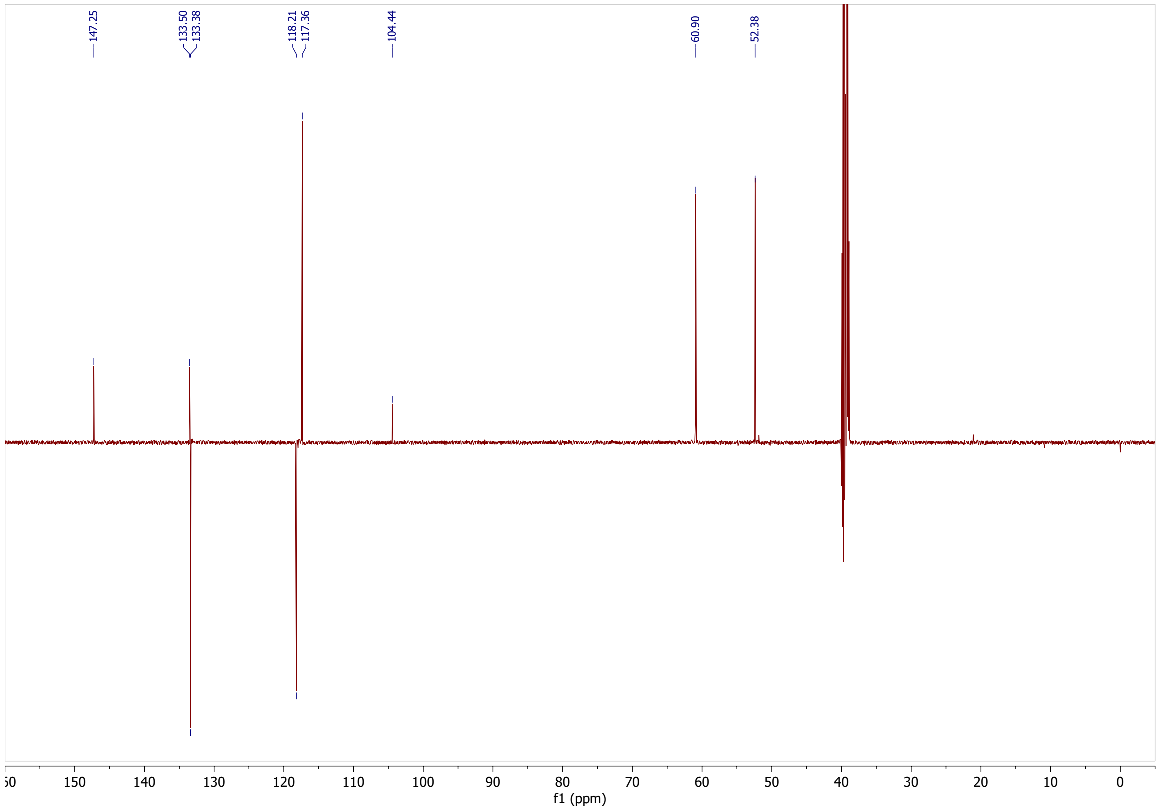

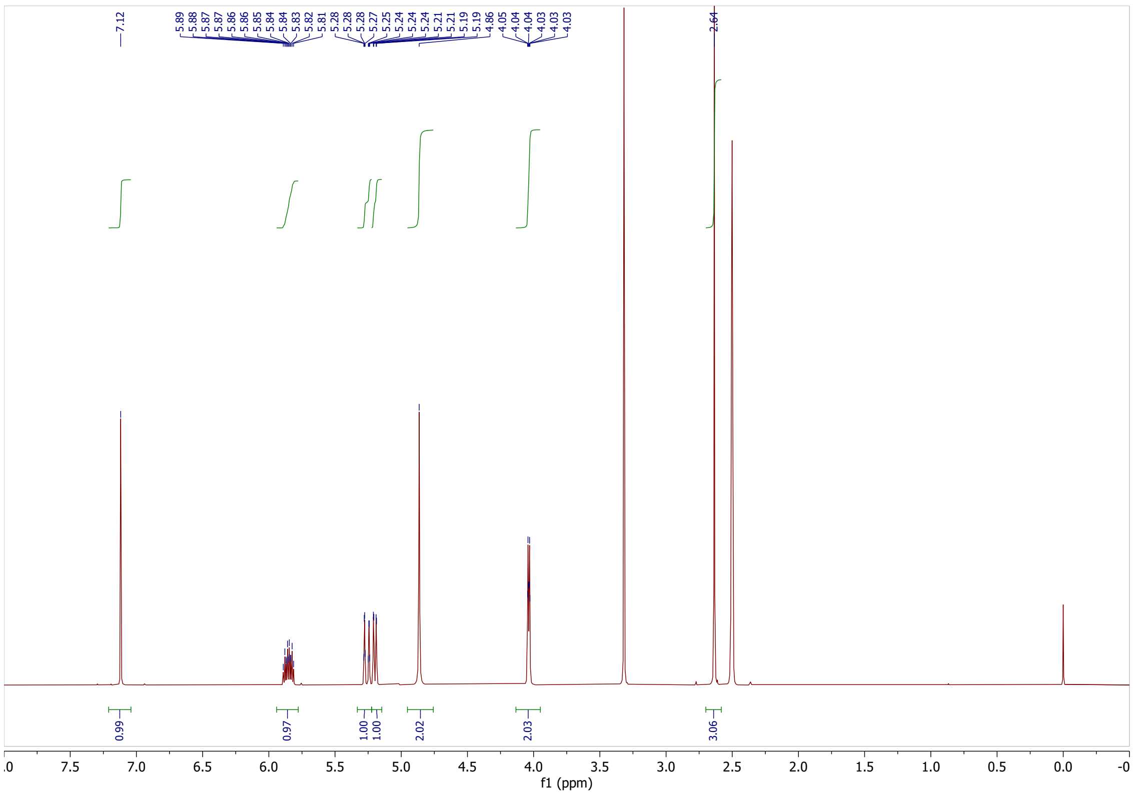
7.12
5.89
5.88
5.87
5.87
5.86
5.86
5.85
5.84
5.84
5.83
5.82
5.81
5.28
5.28
5.28
5.27
5.25
5.24
5.24
5.24
5.21
5.21
5.19
5.19
4.86
4.05
4.04
4.04
4.03
4.03
4.03
2.64
0.99
0.97
1.00
1.00
2.02
2.03
3.06
f1 (ppm)

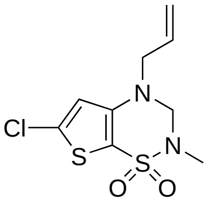
N
Cl
S
N
S
O
O

**28**

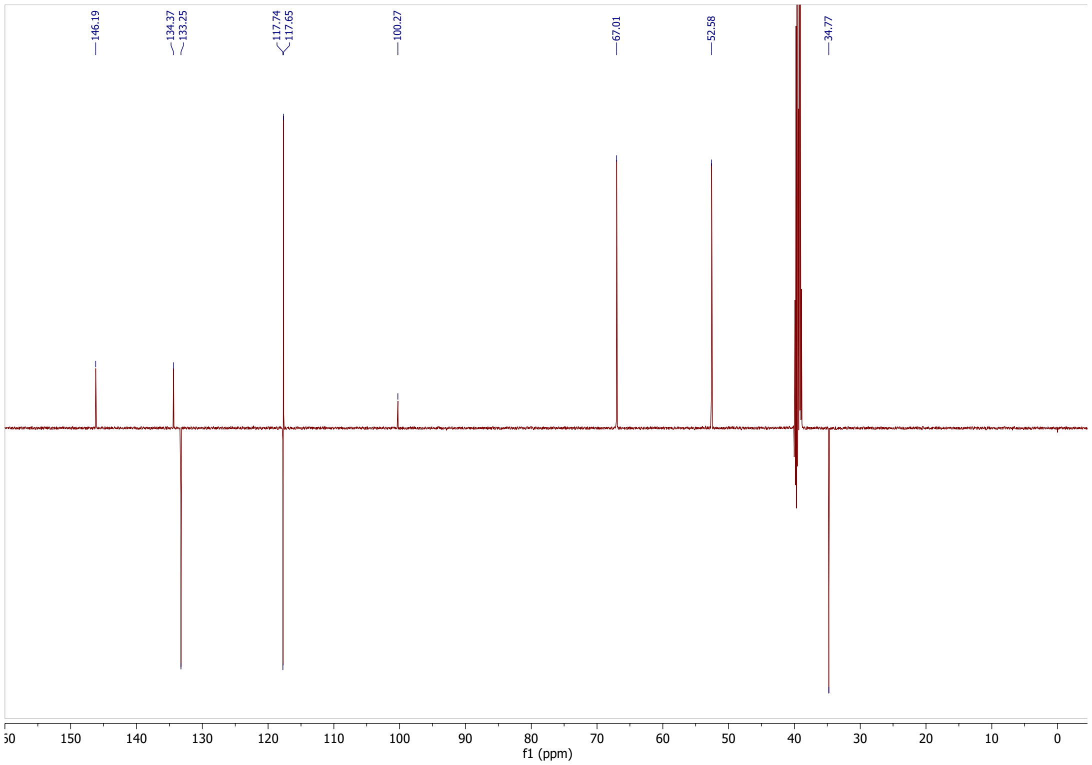
146.19
134.37
133.25
117.74
117.65
100.27
67.01
52.58
34.77
f1 (ppm)

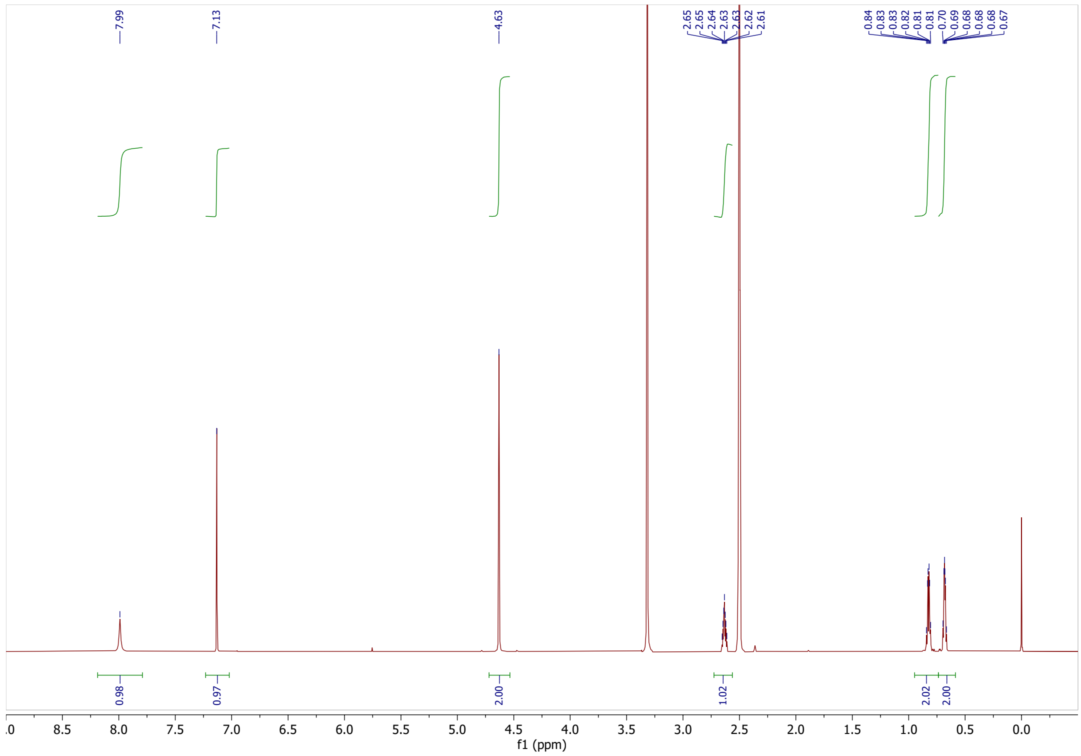
7.99
7.13
4.63
2.65
2.65
2.64
2.63
2.63
2.62
2.61
0.84
0.83
0.83
0.82
0.82
0.81
0.70
0.69
0.68
0.68
0.68
0.67
0.98
0.97
2.00
1.02
2.02
2.00
f1 (ppm)

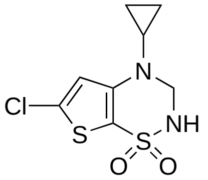
N
Cl
S
NH
S
O
O

**32**

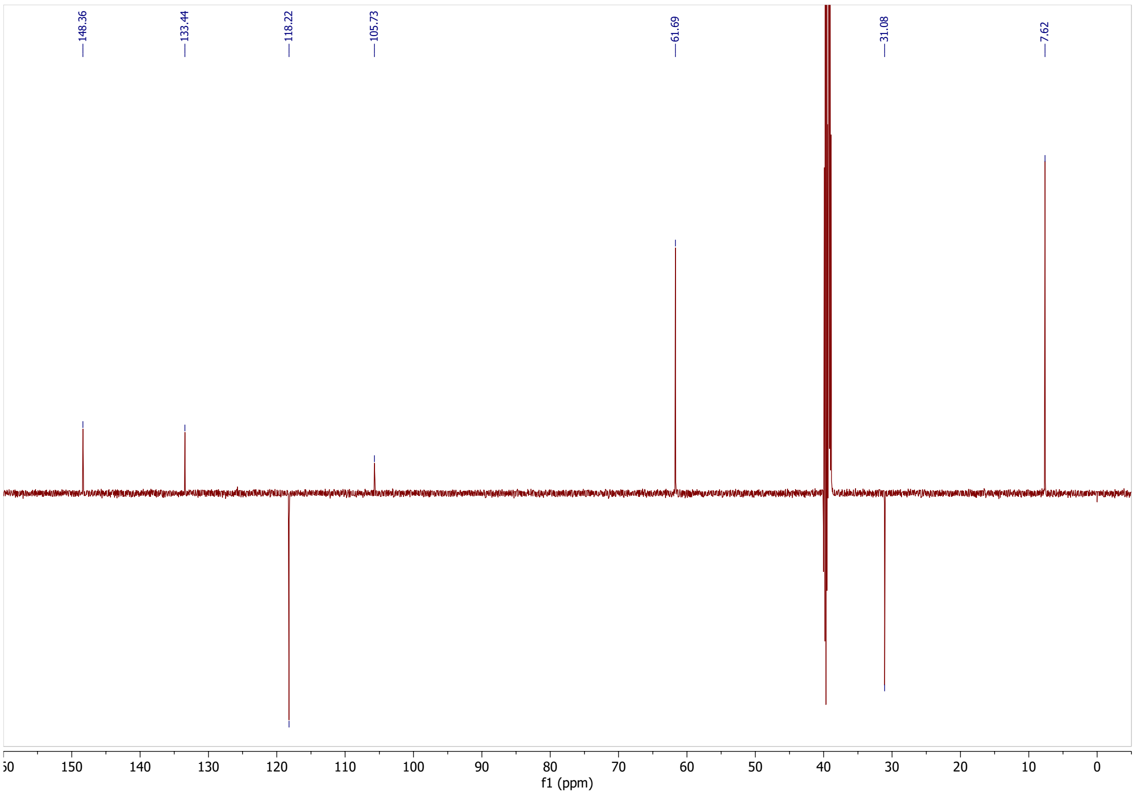
148.36
133.44
118.22
105.73
61.69
31.08
7.62
f1 (ppm)

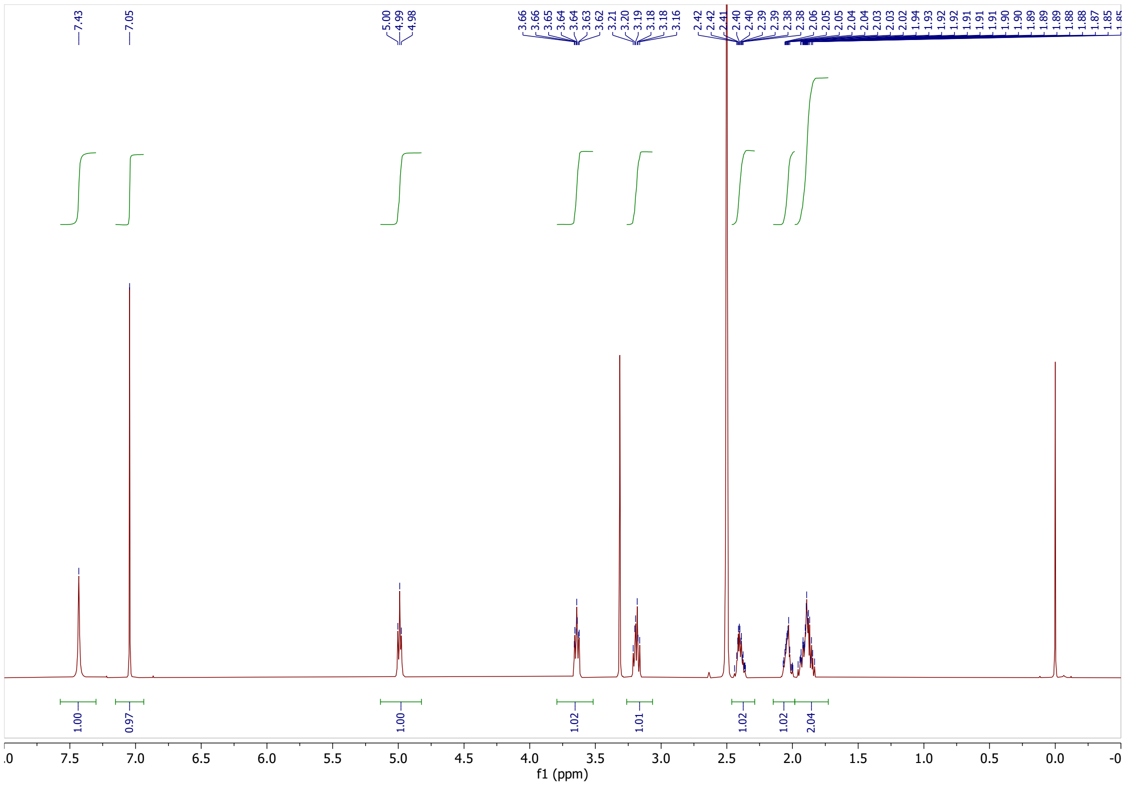
7.43
7.05
5.00
4.99
4.98
1.00
0.97
1.00
1.02
1.01
1.02
1.02
2.04
f1 (ppm)

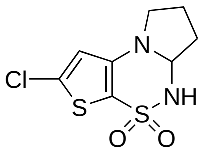
Cl
S
N
NH
S
O
O

**36**

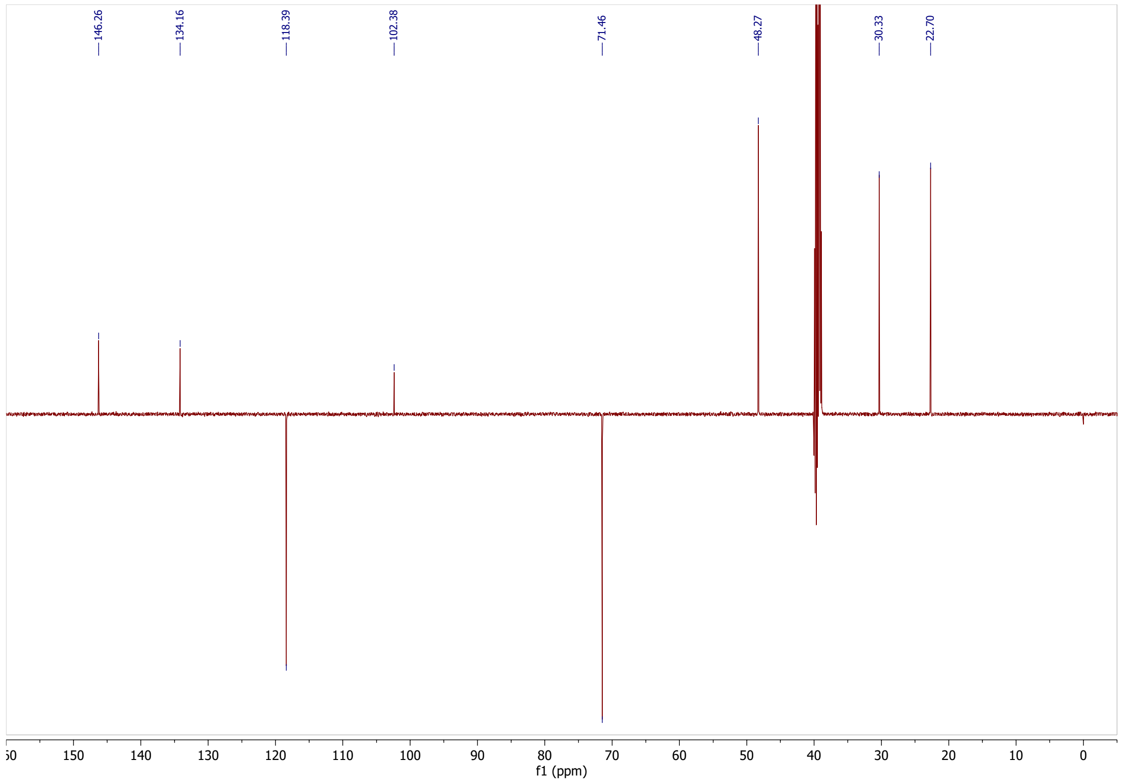
146.26
134.16
118.39
102.38
71.46
48.27
30.33
22.70
f1 (ppm)

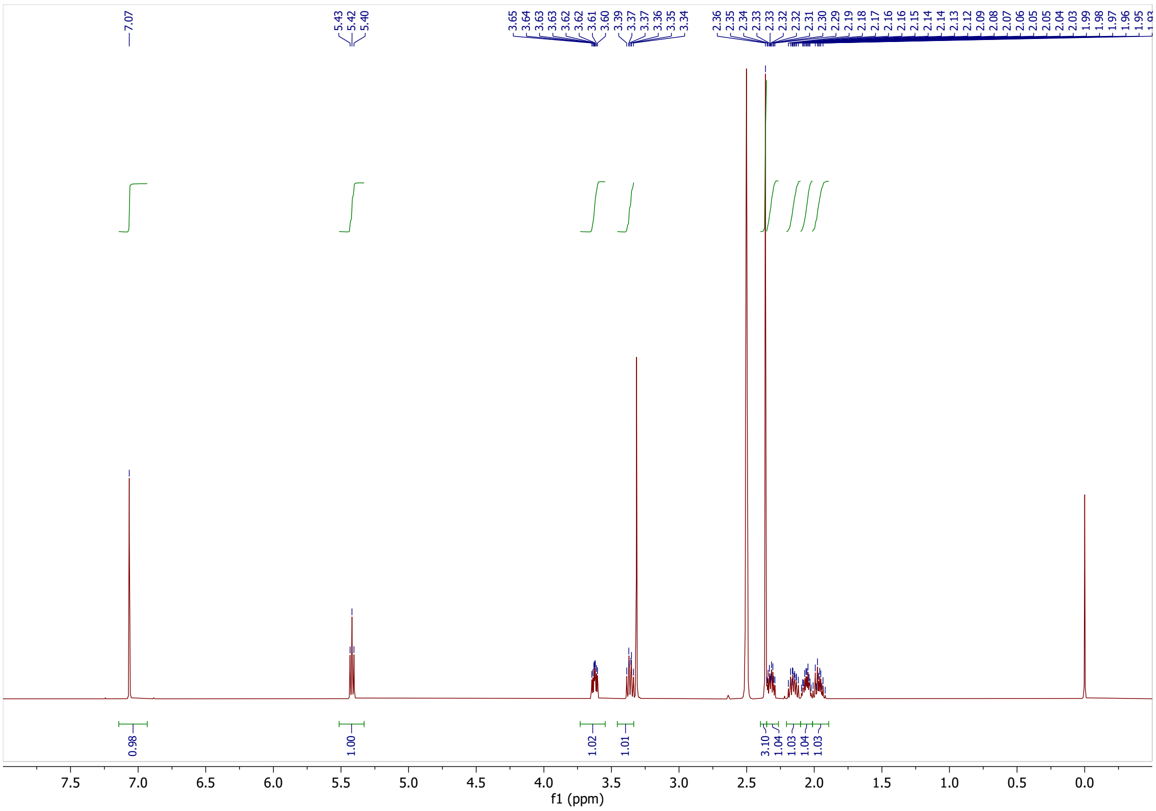
7.07
5.43
5.42
5.40
3.65
3.64
3.63
3.63
3.62
3.62
3.61
3.60
3.39
3.37
3.37
3.36
3.35
3.34
2.36
2.35
2.34
2.33
2.33
2.32
2.32
2.31
2.30
2.29
2.19
2.18
2.17
2.16
2.16
2.15
2.14
2.14
2.13
2.12
2.09
2.08
2.07
2.06
2.05
2.05
2.04
2.03
1.99
1.98
1.97
1.96
1.95
0.98
1.00
1.02
1.01
3.10
1.04
1.03
1.04
1.03
7.5
7.0
6.5
6.0
5.5
5.0
4.5
4.0
3.5
3.0
2.5
2.0
1.5
1.0
0.5
0.0
f1 (ppm)

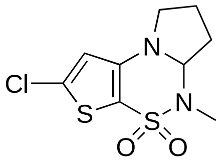
Cl
S
N
N
S
O
O

**37**

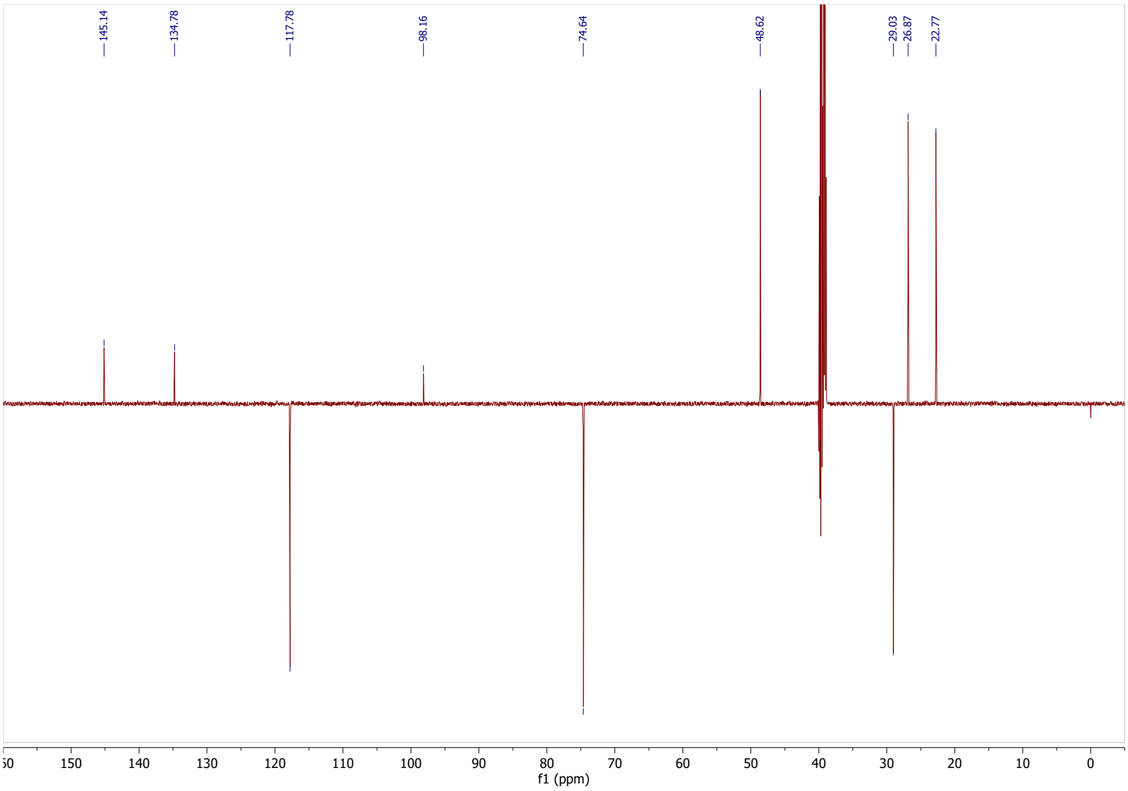
145.14
134.78
117.78
98.16
74.64
48.62
29.03
26.87
22.77
150
140
130
120
110
100
90
80
70
60
50
40
30
20
10
0
f1 (ppm)

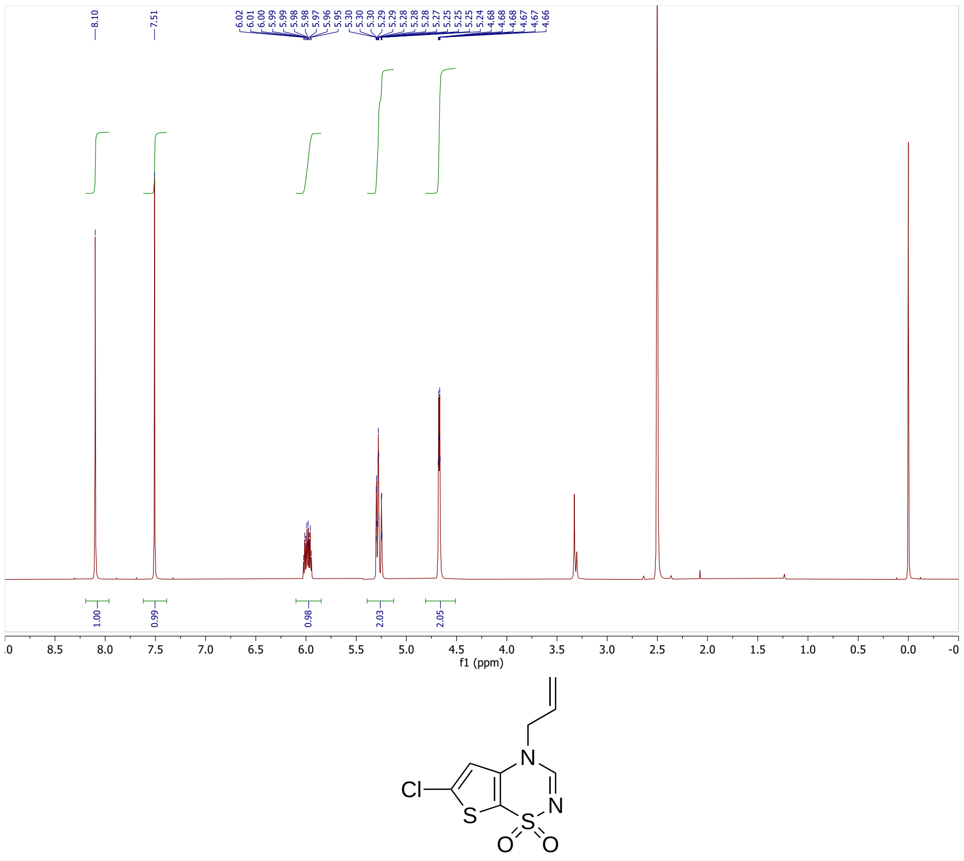
8.10
7.51
1.00
0.99
0.98
2.03
2.05
f1 (ppm)
Cl
S
N
S
O
O
N

**26**

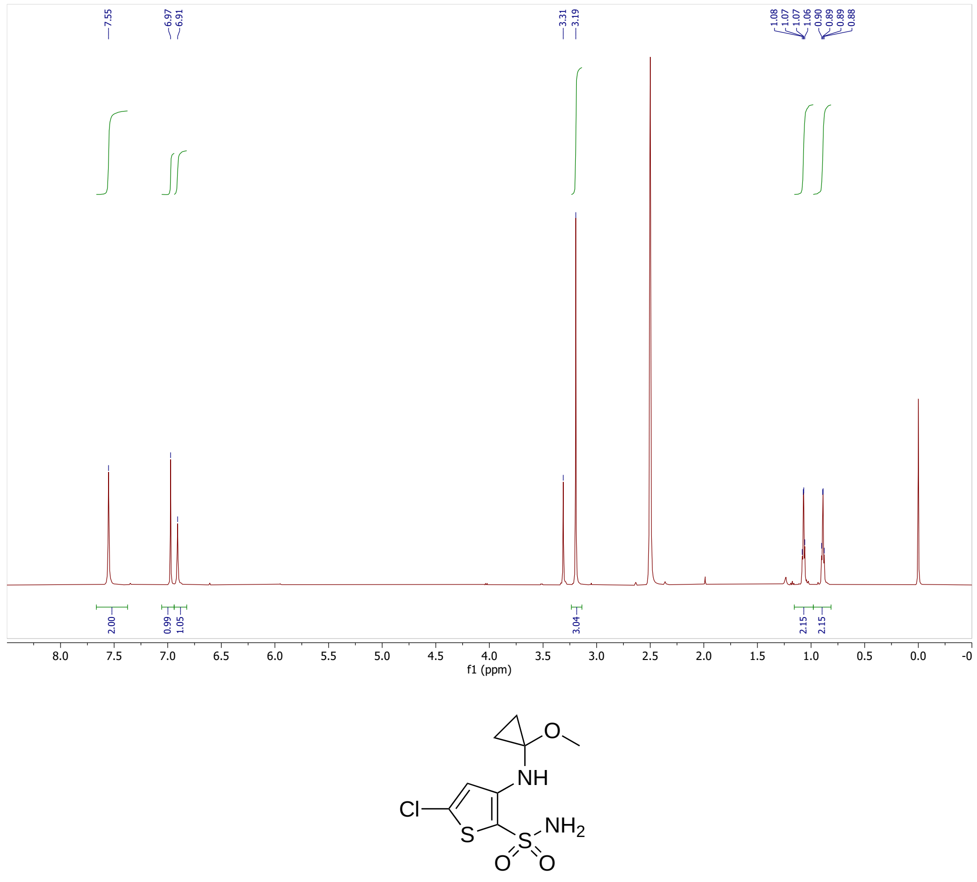
7.55
6.97
6.91
3.31
3.19
2.00
0.99
1.05
3.04
2.15
2.15
f1 (ppm)
O
NH
Cl
S
NH2
S
O
O

**29**

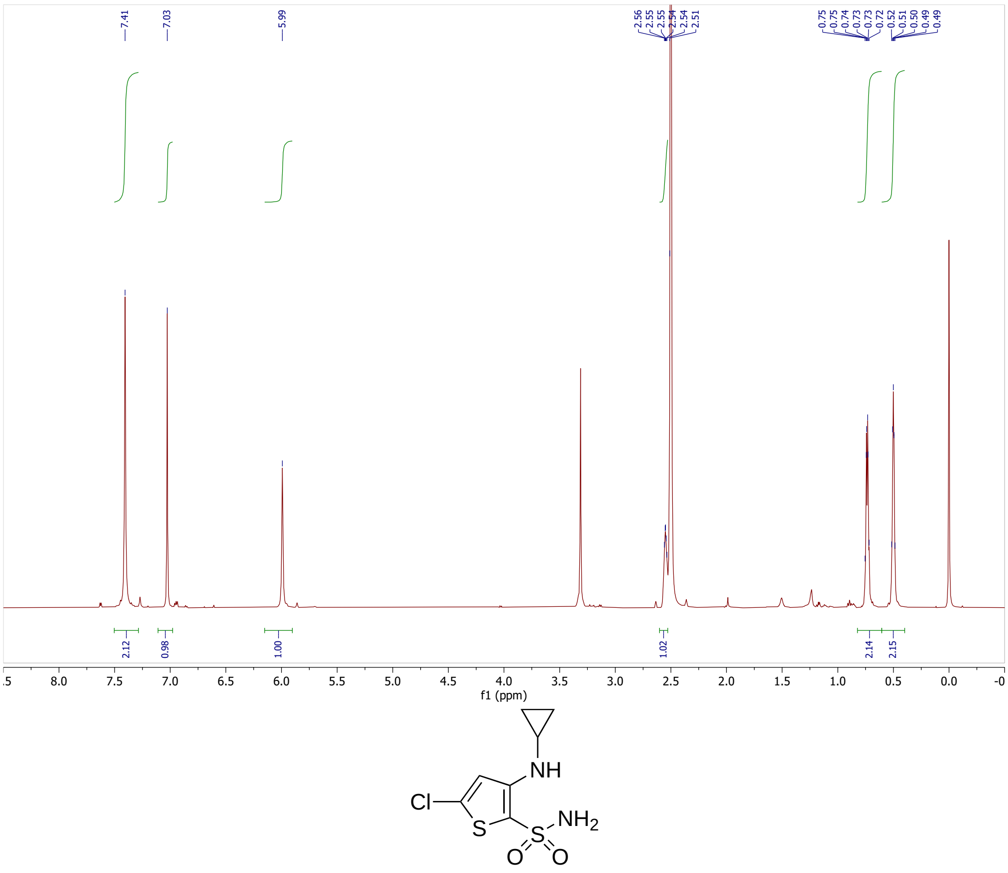
7.41
7.03
5.99
2.56 2.55 2.55 2.54 2.54 2.51
0.75 0.75 0.74 0.73 0.73 0.72 0.52 0.51 0.50 0.49 0.49
2.12
0.98
1.00
1.02
2.14
2.15
f1 (ppm)
NH
Cl
S
$NH_2$
S
O O

**30**

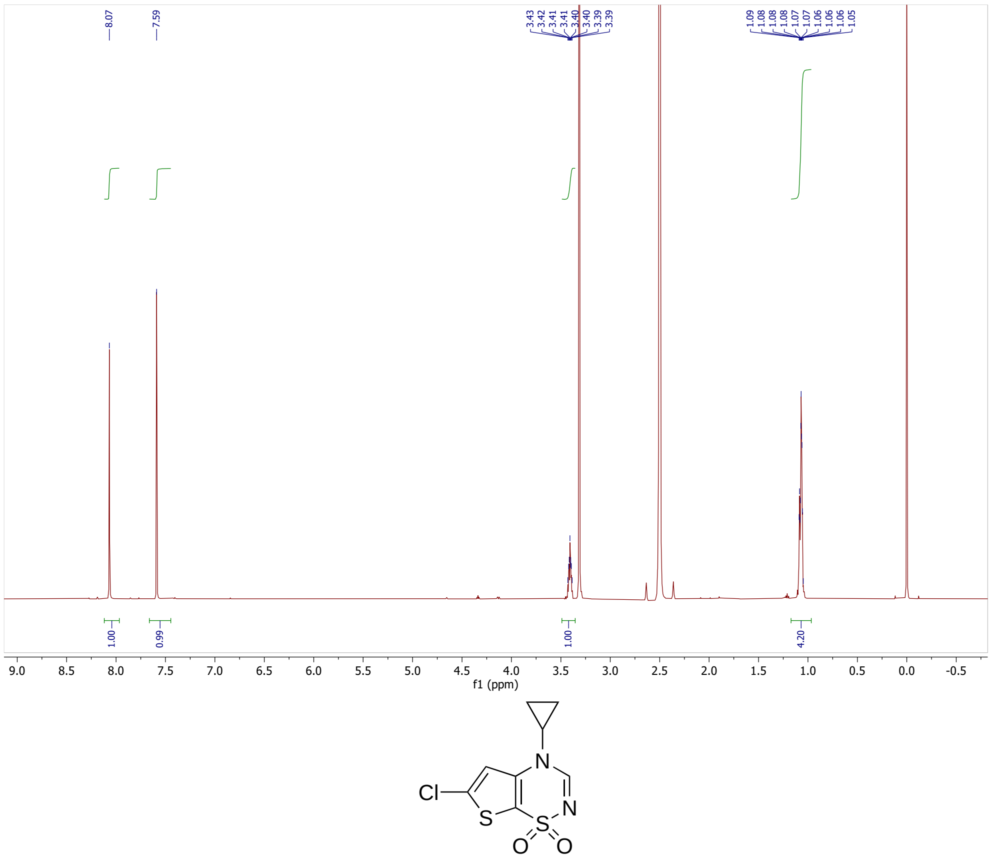
8.07
7.59
3.43 3.42 3.41 3.40 3.40 3.39 3.39
1.09 1.08 1.08 1.08 1.07 1.07 1.06 1.06 1.06 1.05
1.00
0.99
1.00
4.20
f1 (ppm)
N
Cl
N
S
S
O O

**31**

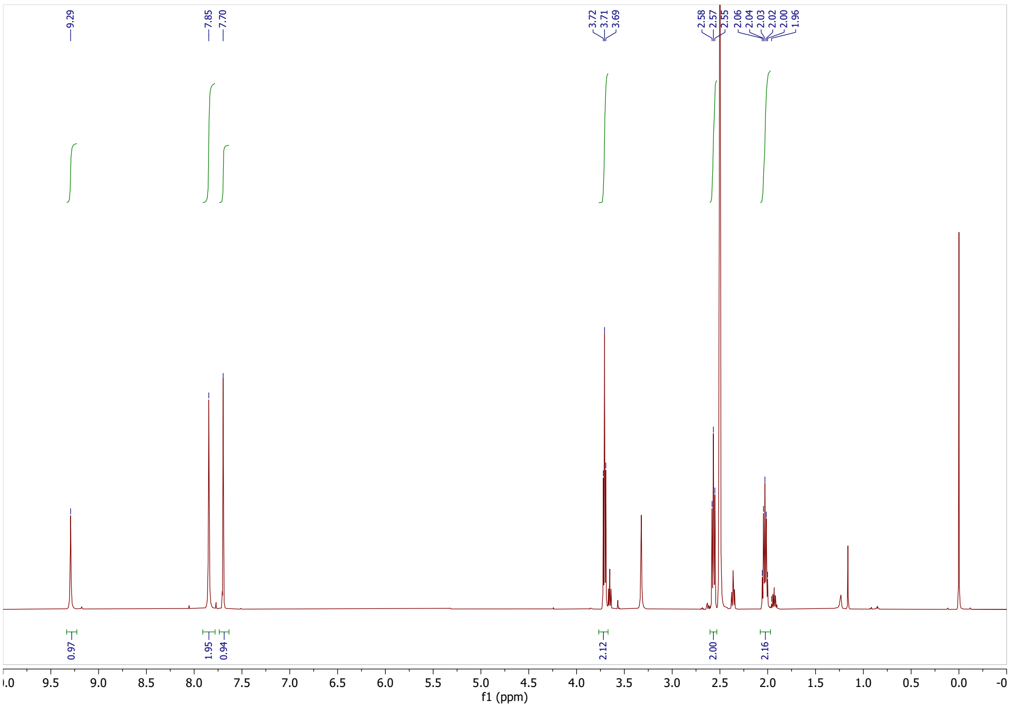
9.29
7.85
7.70
3.72
3.71
3.69
2.58
2.57
2.55
2.06
2.04
2.03
2.02
2.00
1.96
0.97
1.95
0.94
2.12
2.00
2.16
f1 (ppm)

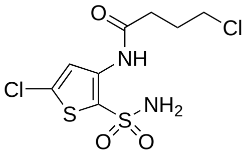
O
NH
Cl
Cl
S
$NH_2$
S
O
O

**33**

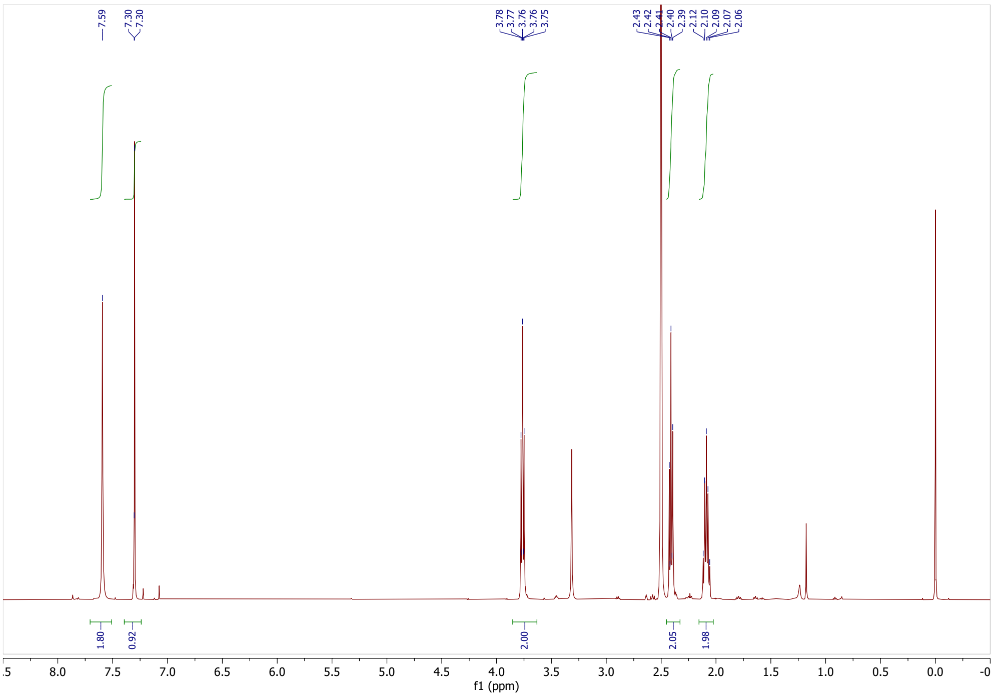
7.59
7.30
7.30
3.78
3.77
3.76
3.76
3.75
2.43
2.42
2.41
2.40
2.39
2.12
2.10
2.09
2.07
2.06
1.80
0.92
2.00
2.05
1.98
f1 (ppm)

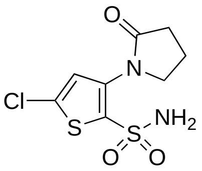
O
N
Cl
S
$NH_2$
S
O
O

**34**

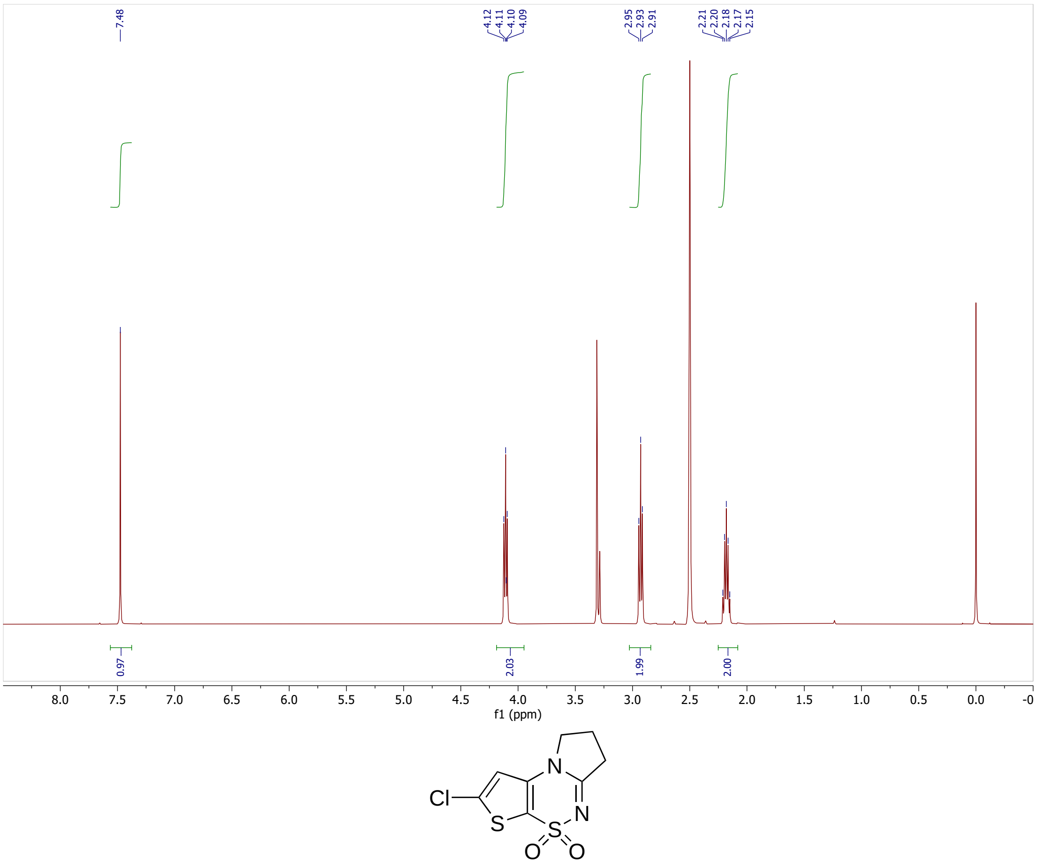
7.48
4.12
4.11
4.10
4.09
2.95
2.93
2.91
2.21
2.20
2.18
2.17
2.15
0.97
2.03
1.99
2.00
8.0 7.5 7.0 6.5 6.0 5.5 5.0 4.5 4.0 3.5 3.0 2.5 2.0 1.5 1.0 0.5 0.0
f1 (ppm)
Cl
S
N
N
S
O
O

**35**
